# Microenvironmental and transcriptional determinants of the robustness of spinal cord regeneration in zebrafish

**DOI:** 10.64898/2026.09.07.749713

**Authors:** Pierre Gillotay, Sushant Bangru, Jingwen Shen, Bethany Moore, Jianhong Ou, Ron Stewart, Kenneth D. Poss

## Abstract

Unlike mammals, adult zebrafish can regenerate spinal cord tissue after complete transection. We report here that a substantial proportion of zebrafish receiving this injury fail to regain locomotor function, and we analyzed this critically. After stratifying large cohorts of injured animals according to functional recovery, we directly compared recovered and non-recovered individuals at matched post-injury stages. We find that permanently paralyzed animals form a tissue bridge across the lesion yet exhibit impaired axonal invasion and regrowth to downstream targets. Integrative single-nucleus profiling, spatial transcriptomics, and regulatory network analyses revealed that regenerative success is characterized by the coordinated establishment of a permissive multicellular regenerative state rather than major differences in cellular composition. Furthermore, we find that regenerative failure involves persistent non-permissive regulatory programs, including extracellular matrix remodeling, impaired adaptive T-cell activation, and reduced axon growth-associated signaling. Gene regulatory network and cellular trajectory analyses identified candidate transcriptional regulators that could not be resolved through differential expression analysis alone. Functional CRISPR F_0_ screening of several of these candidates identified *hoxb5a* as a critical determinant of regenerative outcome. Stable *hoxb5a* mutants exhibited impaired swimming recovery, disrupted bridge formation, and limited axon bridging after spinal cord injury. Together, our findings reveal key differences between success and failure in a model of elevated regenerative capacity, with *hoxb5a* acting as a regulator linking multicellular tissue remodeling to axonal regeneration and functional recovery.

## Introduction

Spinal cord injury (SCI) is a common, devastating event that can cause permanent sensory, motor, and autonomic deficits in humans (^1–3)^. The failure to effectively regenerate spinal cord and other central nervous system (CNS) structures has been attributed to a combination of intrinsic limitations in neuronal growth capacity and the formation of a non-permissive lesion environment characterized by persistent inflammation, inhibitory extracellular matrix deposition, and glial and fibrotic scarring (^4–9)^. Recent advances have substantially expanded our understanding of the cellular basis of spinal cord repair. Functional studies have identified specialized neuronal populations capable of orchestrating recovery after injury (^10^), and large-scale single-cell, multi-omic, and spatial transcriptomic analyses have begun to resolve the dynamic cellular states and molecular programs underlying spinal cord injury and regeneration across species (^11,12^). In parallel, neuromodulation studies have demonstrated that some functional recovery can be achieved through coordinated remodeling of spinal circuits and activation of pro-regenerative transcriptional programs (^13^). Together, this work has diversified the field, emphasizing both identification of pro-regenerative factors as well as pro-regenerative cellular networks.

In contrast to mammals, vertebrate species like zebrafish, lamprey, and salamanders possess a remarkable capacity to regenerate the spinal cord following complete transection. Injured adult animals can restore tissue continuity, re-establish descending and ascending axonal connections, and recover locomotor function within weeks after injury (^14–17)^. This regenerative response is supported by coordinated cellular processes that include extracellular matrix (ECM) remodeling, revascularization, dynamic immune cell responses, ependymo-radial glial proliferation and neurogenic differentiation, followed by axonal regrowth across the lesion site (^14,15,18–26)^. Following spinal cord injury in zebrafish, fibroblast-like stromal cells participate in ECM remodeling and production of molecular cues that influence tissue repair (^24,27,28^). At the same time, innate and adaptive immune populations actively shape the pro-regenerative microenvironment through cytokine production, debris clearance, and regulation of cellular behavior within the lesion site (^23,25,26^). In defining events, radial glia contribute to the formation of a permissive tissue bridge that supports axon regeneration (^20,29^).

Comparisons between species with different regenerative capacities have generated hypotheses on what may distinguish outcomes. While informative, massive differences in injury responses, tissue architecture, evolutionary history, and developmental programs can challenge analyses and conclusions (^30–32)^. By contrast, natural variation between strains or populations, and even individuals, is much more limited and can more readily be linked with differences in cellular states and molecular programs of regeneration (^33,34^). Dissecting the sources of such variation could reveal critical determinants of spinal cord regeneration. Here, we find that an unexpectedly high proportion (30%) of adult zebrafish fail to recover locomotor function following complete spinal cord transection. We combined behavioral, transcriptomic, and genetic analyses to show that successful recovery depends on establishing a coordinated pro-regenerative cellular state, whereas non-recovered animals remain trapped in non-permissive programs that limit axon crossing despite formation of a tissue bridge across the injury site. Bioinformatic and molecular genetic approaches identify Hoxb5a as a transcription factor critical to regenerative success, with mutant animals showing impaired recovery, reduced bridge formation, and limited axonal regrowth after spinal cord injury.

## Results

### A large, reproducible subset of zebrafish does not reverse paralysis after spinal cord transection

In examining spinal cord regeneration in a large group of adult zebrafish prepared for transcriptomic analysis, we recognized a relatively high proportion showed poor swimming ability at 6-8 weeks post resection. While we had anecdotally observed the lack of functional recovery in the past, to our knowledge regenerative failures in zebrafish like these have not previously been emphasized or analyzed critically. To characterize this variability in outcome, we performed complete spinal cord transections in over 250 animals aged 4–6 months of the Ekkwill strain and longitudinally assessed locomotor recovery using a swim performance assay (^16,17^). At 3 days post-injury (dpi), animals were screened to confirm complete paralysis, ensuring consistency of lesion severity across the experimental cohort. Animals retaining detectable swimming ability at this stage were excluded from further analysis (see Methods).

Over a six-week recovery period, injured adult animals diverged into two reproducible functional outcomes. While most regained the ability to sustain swimming against increasing water flow, a notable subset (∼28%, 93/254) remained unable to tolerate minimal flow conditions (7 cm/s) even at 6 weeks post-injury (wpi) (“non-recovered”; Fig. 1a-c). This divergence was reproducible across biological replicates and independent of sex, and it was observed with different surgeons (Fig. S1). This observation is also consistent with assessment of small cohorts in previous reports from our group and others (^12,29,35–37)^. Uninjured and sham-operated animals consistently outperformed injured groups (recovered and non-recovered), confirming that locomotor deficits were specific to spinal cord transection. To independently test these observations, we quantified spontaneous locomotor activity in a free-swimming environment using an enclosed behavioral tracking system. At 2 wpi, recovered and non-recovered animals displayed comparable total swimming distances despite clear differences in swim tunnel performance (Fig. 1d, e). By 4 wpi, however, recovered animals exhibited significantly increased locomotor activity relative to non-recovered animals (Fig. 1d, e), a difference that persisted at 6 wpi (Fig. 1d, e).

**Figure 1.**
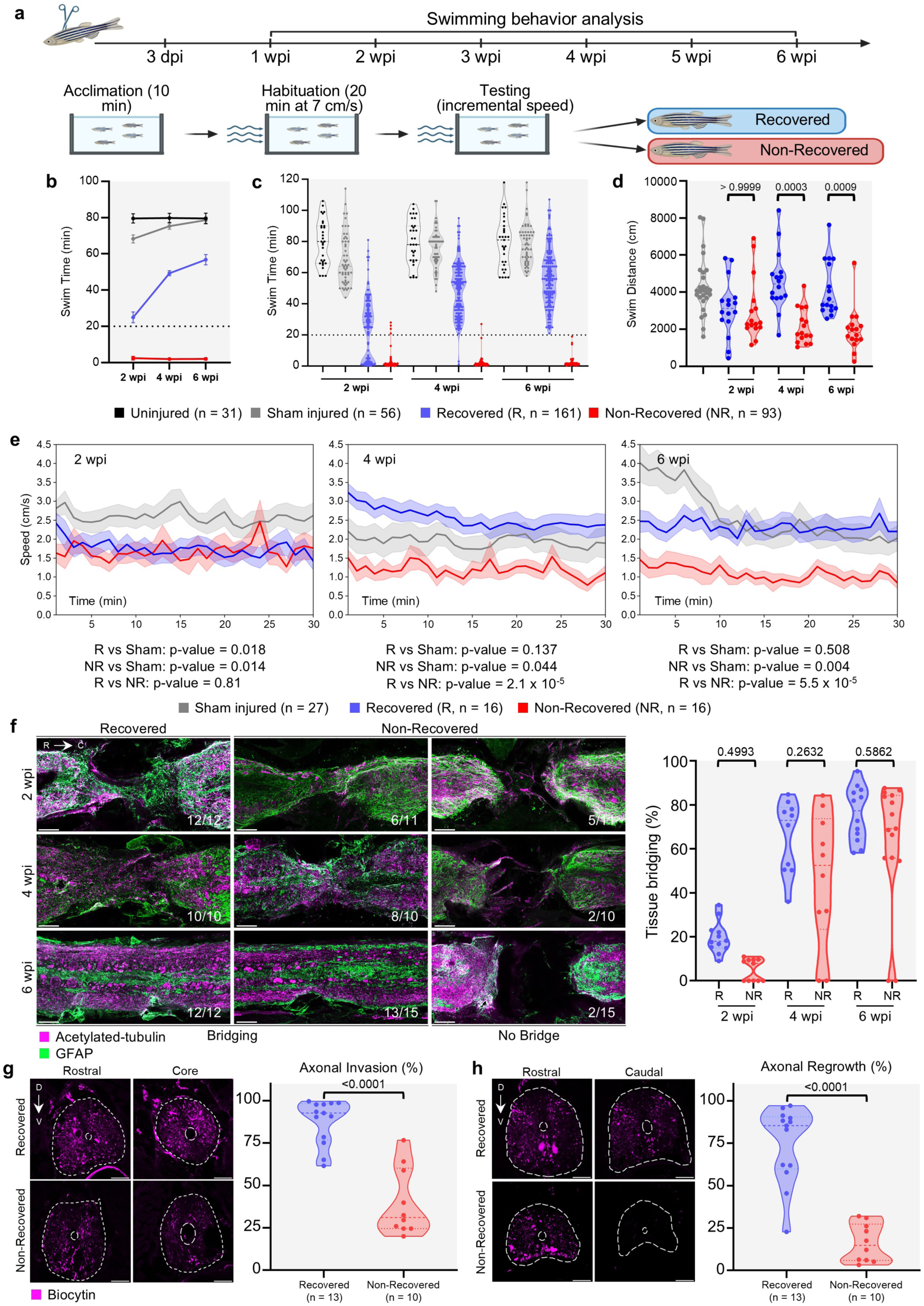
A high incidence of regenerative failure after spinal cord injury in zebrafish. **A)** Experimental pipeline to measure swim function recovery at 2, 4 and 6 weeks post-injury (wpi) **B, C)** Swim capacity of uninjured (black), sham-injured (gray), recovered (blue) and non- recovered animals (red) at increasing current over a 6-week follow-up period after spinal cord transection. Data are shown as line graph **(B)** at 2, 4 and 6 wpi and violin plot **(C)** at the similar timepoint. Each dot represents an individual fish. Dashed line indicates the 20 min acclimatation period and serves as a threshold to identify non-recovered animals. Error bars in **(B)** indicate SEM. **D)** Free swim capacity measurement of sham-injured (gray, n = 27), recovered (blue, n = 16) and non-recovered (red, n=16) animals at the indicated timepoint. A two-tailed Mann–Whitney test was used for comparisons of the recovered and non-recovered animal swim times at the specified timepoint. **E)** Time-resolved swim speed profiles of sham-injured (gray, n = 27), recovered (R, blue, n = 16), and non-recovered (NR, red, n = 16) zebrafish at 2, 4, and 6 weeks post-injury, respectively. Fish were monitored for 30 min in a free-swim assay, and average swim speed was quantified in 1-min bins. Data are presented as mean ± SEM. Statistical analysis performed on average speed comparison using Welch t-test analysis. **F)** Analysis of tissue bridging following spinal cord transection in Recovered and Non-Recovered animals. Representative longitudinal tissue sections indicating the glial marker GFAP (green) and axonal marker acetylated α-tubulin (magenta) in recovered and non-recovered animals at 2, 4 and 6 wpi are shown. n = 10-15. Quantification of tissue bridge thickness is shown on the right side. A two-tailed Mann–Whitney test was used for comparisons. Scale bars, 100 µm. **G)** Analysis of axonal invasion after anterograde axon tracing at 4 wpi. (Left) Representative transverse sections rostral to and at the injury site are shown with biocytin-labelled axons (magenta) and white dashed line highlighting spinal cord and central canal boundaries. (Right) Quantification. A two-tailed Mann–Whitney test was used for comparisons. n = 10-13. Scale bars, 50 µm. **H)** Analysis of axonal regrowth after anterograde axon tracing at 4 wpi. (Left) Representative transverse sections rostral and caudal to the injury site are shown with biocytin-labelled axons (magenta) and white dashed line highlighting spinal cord and central canal boundaries. (Right) Quantification. A two-tailed Mann–Whitney test was used for comparisons. n = 10-13. Scale bars, 50 µm.

### Regenerative failures display full-thickness tissue bridges with limited axon crossing

To determine whether differences in functional recovery reflect underlying structural differences, we next examined tissue morphology and axonal regrowth following injury. Histological analysis revealed that both recovered and non-recovered animals formed tissue bridges connecting the rostral and caudal spinal cord stumps. Quantification of bridge thickness by simultaneous assessment of axonal and glial markers revealed no measurable differences between groups at 2, 4, or 6 wpi (Fig. 1f), indicating that gross anatomical continuity alone is insufficient to restore locomotor function.

To directly assess axonal regeneration, we performed retrograde tracing using fluorescent biocytin. At 4 wpi, animals previously classified as recovered or non-recovered underwent a secondary lesion rostral to the original injury site followed by localized application of fluorescent biocytin. Tissue was collected 24 hours later and sectioned to quantify axonal labeling anterior to, within, and posterior to the lesion bridge. Recovered animals displayed robust axonal invasion into the lesion site, with an average invasion index of ∼95%, whereas axonal invasion was markedly reduced in non-recovered animals, with an average invasion index of ∼30% (Fig. 1h). Consistent with this initial difference, recovered animals showed extensive axonal extension beyond the lesion region, with regenerating axons detected up to 200 µm caudal to the injury site and an average regrowth index approaching 80% (Fig. 1g). In contrast, non-recovered animals exhibited minimal axonal extension beyond the lesion bridge, with regrowth indices averaging ∼15% (Fig. 1g). Together, these findings demonstrate that regenerative failure is not associated with impaired tissue bridging at a gross level, but rather with failure of regenerating axons to effectively extend across the lesion site to posterior targets.

### Cell state transitions associated with successful spinal cord regeneration

To define cellular and molecular programs associated with divergent regenerative outcomes, we performed single-nucleus RNA sequencing (snRNA-seq) of adult zebrafish spinal cord tissue following complete spinal cord transection. Tissue was isolated from a ∼4 mm spinal cord segment centered on the lesion site, encompassing both rostral and caudal tissue adjacent to the injury. Samples included sham-injured controls together with injured animals collected at 1, 2, 4, and 6 wpi. Animals collected at 2, 4, and 6 wpi were first classified as recovered or non-recovered using the swim performance assay, whereas animals collected at 1 wpi were analyzed as a single injury group prior to the emergence of stable functional divergence (Fig. 2a).

**Figure 2.**
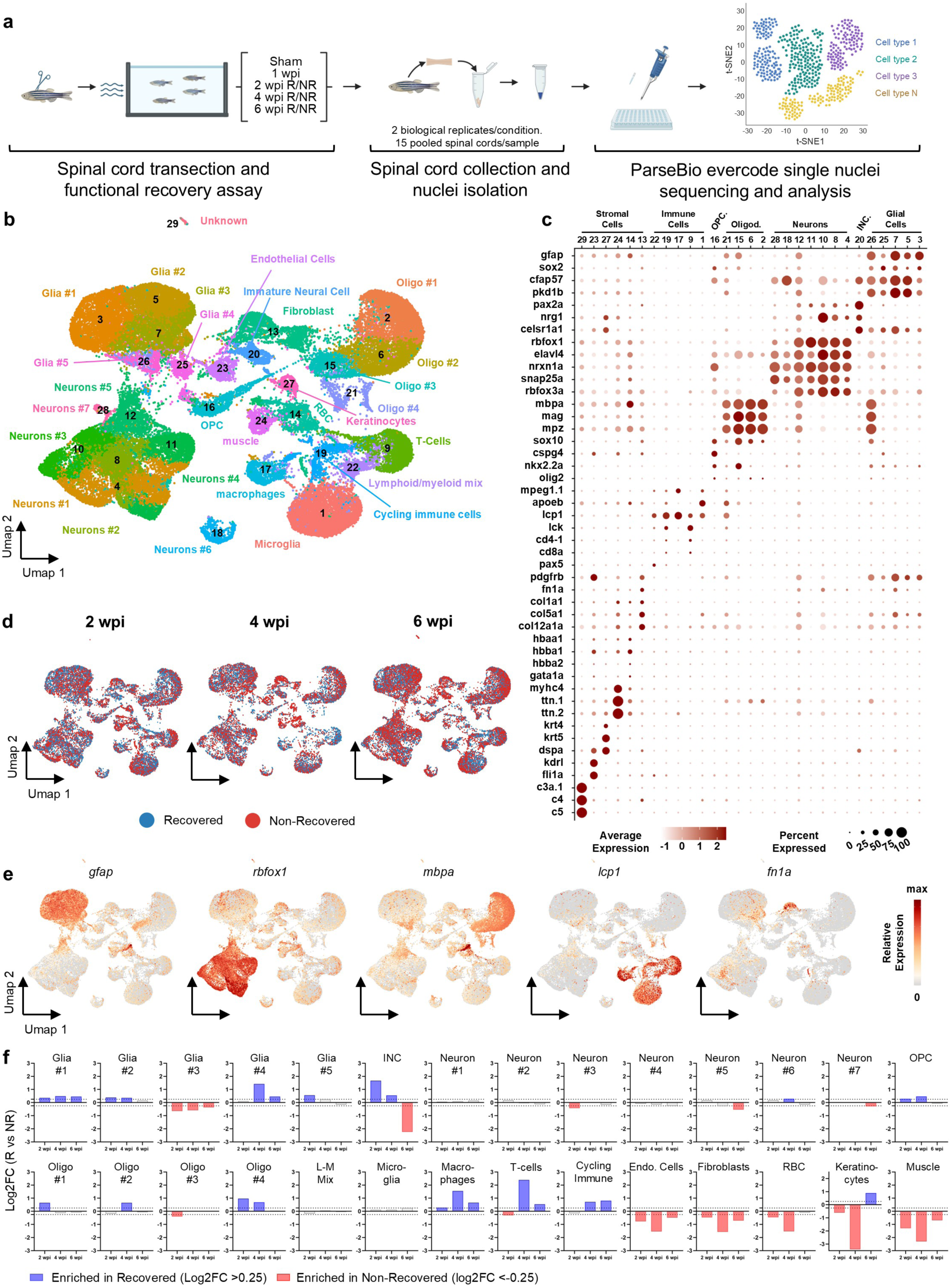
Single-nuclei transcriptomics reveals progressive cellular divergence between recovered and non-recovered animals. **A)** Experimental pipeline to generate a single-nuclei atlas of successful and failed spinal cord regeneration. Spinal cord tissue collection, nuclei isolation and sequencing were performed on wild-type animals at 1, 2, 4 and 6 wpi. Sham-injured spinal cord nuclei were used as controls. Samples at 2, 4 and 6 wpi were collected from recovered and non-recovered animals separately. Two biological replicates were used at each time point and condition, and tissues from 15 animals were pooled into a single biological replicate. Parse Bioscience Evercode Whole Transcriptomic 100k kit was used. **B)** Merged Umap representation of the complete single nuclei dataset. Eight conditions (sham, 1 wpi, 2 wpi – NR, 2 wpi – R, 4 wpi – NR, 4 wpi – R, 6 wpi – NR and 6 wpi -- R) and 2 biological replicates/condition clustered into 29 transcriptionally distinct cell populations. **C)** Marker gene expression in the major cell types identified in **(B)**. Dot plot shows enriched canonical markers and their respective cell types. Dot colors and diameters represent average gene expression and percent cells with at least one UMI detected per gene, respectively. **D)** Umap plots representing the overlap of recovered and non-recovered samples at given timepoints. Blue dots represent recovery-associated cells and red dots represent non-recovery-associated cells **E)** Feature plot showing distribution of canonical cell marker expression representing major cell-types identified in the dataset. **F)** Cluster enrichment comparison between samples from recovered and non-recovered animals at 2, 4 and 6 wpi. Cell proportions were normalized to the total number of cells present at that time point before representation as a log2FC bar plot. For each cluster, timepoints showing log2FC >0.25 are highlighted in blue and log2 FC <-0.25 are highlighted in red.

Following quality-control filtering, 61,485 nuclei were retained for downstream analysis. Unsupervised clustering resolved 29 transcriptionally distinct populations representing the major neural, glial, stromal, vascular and immune cell types expected within the adult zebrafish spinal cord (Fig. 2b). Cell identities were assigned based on established marker genes and transcriptional signatures (Fig. 2c, e; Table S1), identifying multiple glial, neuronal, oligodendrocyte, immune, fibroblast/perivascular fibroblast, endothelial and epithelial populations. In addition, a distinct oligodendrocyte progenitor cell (OPC) population and a small proliferating immune cell population were identified, consistent with the cellular diversity previously reported in the adult zebrafish spinal cord (^12,17,29^). We also detected a minor muscle cell population and an erythrocyte population, likely reflecting contamination during tissue dissection and nuclei isolation.

To determine whether regenerative success and failure were associated with changes in cellular composition, we quantified the relative abundance of each cellular population across regeneration outcomes and time points. Whereas most neuronal and glial populations remained broadly stable throughout regeneration, several stromal and immune-associated populations displayed pronounced condition-dependent remodeling during the regenerative process (Fig. 2d, f). Importantly, these differences progressively emerged between 4 and 6 wpi, coinciding with the establishment of stable functional divergence between recovered and non-recovered animals. Among all identified populations, fibroblasts (Cluster 13) displayed one of the strongest enrichments in non-recovered animals, peaking at 6.1% of total nuclei at 4 wpi in non-recovered animals compared to 2.4% in their recovered siblings, suggesting persistent stromal activation during regenerative failure (Fig. 2f). In contrast, a T-cell-associated population (Cluster 9) was comparatively enriched in recovered animals at 4 wpi (5.4% in recovered vs 1.1% in non-recovered, Fig. 2f). Interestingly, the immature neural cell populations, represented in Cluster 20, also progressively accumulated in non-recovered animals at later stages of regeneration (0.7% in recovered vs 3.2% in non-recovered at 6 wpi, Fig. 2f), suggesting the persistence of unresolved cellular remodeling states during failed spinal cord repair. Together, these findings indicate that regenerative success and failure are associated with selective remodeling of specific stromal, immune, and morphogenetic cellular constituents rather than broad alterations in overall spinal cord cellular composition.

### Targeted spatial transcriptomics reveals a localized pro-fibrotic compartment

To determine whether the transcriptional differences identified by single-nucleus RNA sequencing were accompanied by spatial remodeling of the regenerating spinal cord, we performed targeted spatial transcriptomics using the Xenium platform (10X Genomics). Longitudinal spinal cord sections spanning the lesion site were collected at 1, 2, 4 and 6 wpi, including recovered and non-recovered animals from 2 wpi onward (Fig. 3a). A custom 100-gene panel was designed using cell identity markers identified by our single-nucleus dataset together with genes representing major biological processes involved in spinal cord regeneration (Table S2).

**Figure 3.**
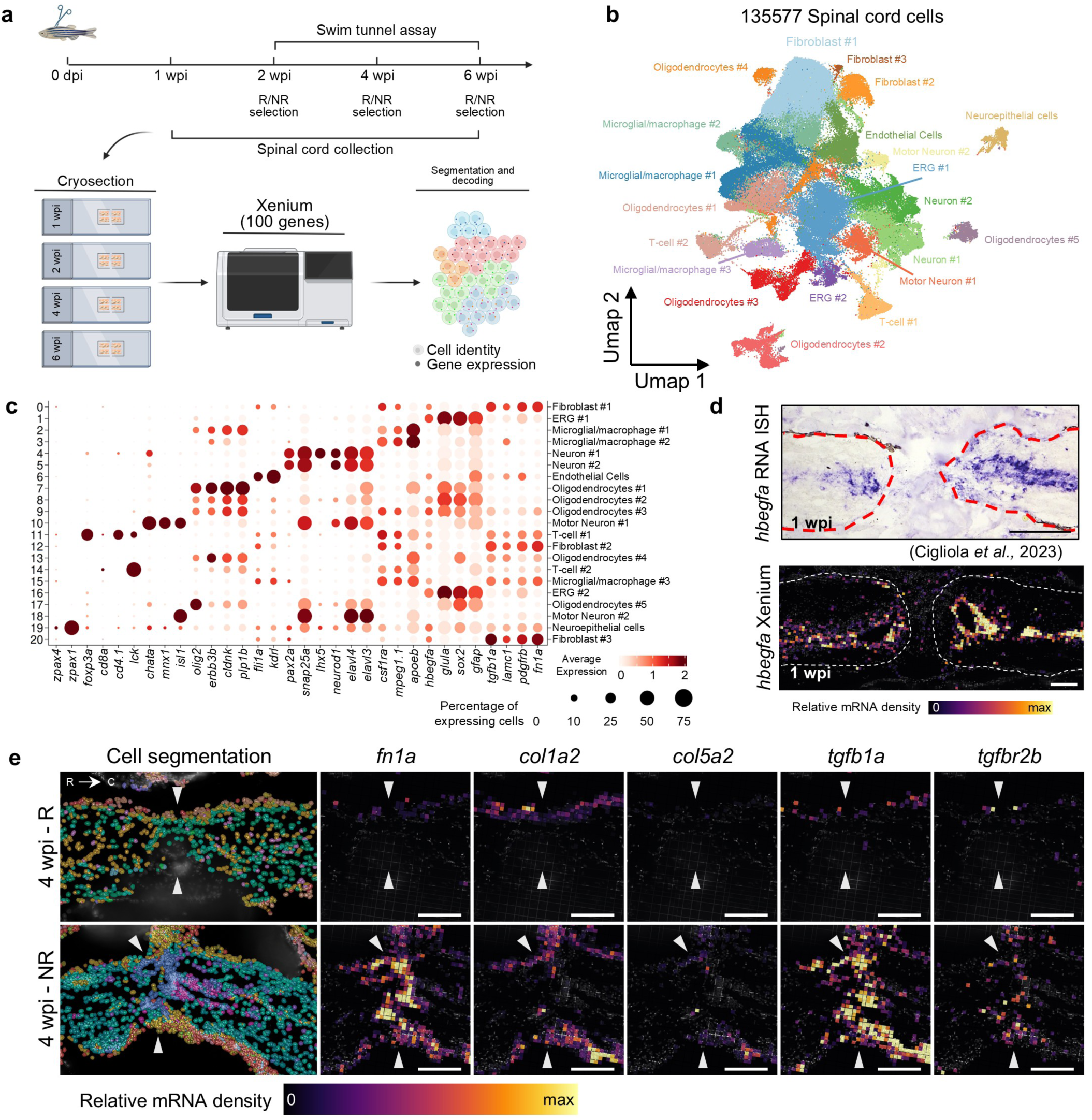
Spatial transcriptomics reveals fibrosis-associated remodeling during failed spinal cord regeneration. **A)** Experimental pipeline followed to generate a spatial transcriptomic dataset of successful and failed spinal cord regeneration using the Xenium platform from 10X Genomics. **B)** Merged Umap representation of spinal cord tissue sections used in spatial transcriptomic experiments. Seven conditions (1 wpi (n = 4), 2 wpi – NR (n = 7), 2 wpi – R (n = 4), 4 wpi – NR (n = 4), 4 wpi – R (n = 6), 6 wpi – NR (n = 8), and 6 wpi – R (n = 4)) were used, and multiple biological replicates/condition clustered into 21 transcriptionally distinct cell populations. **C)** Marker gene expression in the major cell types identified in **(B)**. Dot plot shows enriched canonical markers and their respective cell types. Dot colors and diameters represent average gene expression and percent cells with at least one UMI detected per gene, respectively. **D)** Comparison of *hb-egfa* expression pattern on longitudinal section of adult zebrafish spinal cord at 1 wpi using colorimetric in situ hybridization (^17^) and Xenium spatial transcriptomic data (Bottom, data are represented has an *hb-egfa* relative RNA density map, bins of 10*10 µm). Scale bars: 200 µm (top), 100 µm (bottom) **E)** Density map expression of fibrosis markers in longitudinal adult zebrafish spinal cord tissue sections at 4 wpi during successful (4 wpi-R, top) or failing (4 wpi-NR, bottom) regeneration. From left to right: cell segmentation map, *fn1a*, *col1a2*, *col5a2*, *tgfb1a* and *tgfbr2b* expression patterns. Results are shown as relative RNA density heatmap (black to yellow) on 10*10 µm bins. White arrowheads indicate location of spinal cord transection. Scale bars, 100 µm.

Following quality control, ∼700,000 cells were segmented across all conditions and clustered into 21 transcriptionally distinct clusters representing 11 major cell populations (Fig. 3b, c, Fig. S3). Spatial gene expression recapitulated known anatomical expression domains, supporting the quality of the dataset. For example, the injury-induced growth factor *hb-egfa* reproduced its previously reported expression pattern (Fig. 3d), while canonical cell identity markers localized to their expected anatomical compartments. *gfap* expression outlined the ependymo-radial glial layer surrounding the central canal and extending through the injury bridge, *mbpa* labeled oligodendrocytes throughout the spinal cord parenchyma, *pdgfrb* marked fibroblasts enriched within the lesion core, and *apoeb* highlighted microglia dispersed throughout uninjured tissue but accumulating at the injury site during the early regenerative response (Fig. S3).

To determine if spatial organization differed between successful and failed regeneration, we compared anatomical distributions of cellular markers between recovered and non-recovered animals. Among major groups, ECM remodeling genes showed strong spatial divergence. At 4 wpi, expression of the fibrosis-associated genes *fn1a, col1a2, tgfb1a* and *tgfbr2b* became highly concentrated within the lesion core of non-recovered animals but remained low or largely absent from the corresponding region of recovered siblings (Fig. 3e). In contrast, expression of *col5a2* remained minimal in both regeneration outcomes, indicating that ECM remodeling during regenerative failure is selective rather than reflecting a generalized increase in collagen deposition. We infer from this analysis that regenerative failure is associated with the emergence of a spatially restricted pro-fibrotic lesion microenvironment that is not present during successful regeneration. Together with our single-nucleus analyses, these results indicate that regenerative failure occurs not only due to altered cellular transcriptional states but also due to their spatial organization within the injury site.

### Immune and stromal compartment diversification during regeneration

To gain deeper insight into the cellular mechanisms underlying divergent regenerative outcomes, we next examined the immune and stromal compartments identified in our snRNA-seq dataset. The immune compartment encompassed microglia, macrophages, T cells, B cells, neutrophils, and smaller proliferating immune populations, whereas the stromal compartment was primarily represented by fibroblast/perivascular fibroblast populations. Among these populations, T cells (Cluster 9) and fibroblasts (Cluster 13) showed particularly strong associations with regenerative outcome and were therefore examined in greater detail. We noted that T-cell transcriptional programs between recovered and non-recovered animals revealed limited differences at 2 and 6 wpi, with no significantly enriched biological processes detected at either timepoint (Table S3). In contrast, substantial divergence emerged at 4 wpi, with differential enrichment of gene ontology (GO) terms predominantly associated with positive regulation of transcription, RNA biosynthesis, NF-κB signaling, leukocyte differentiation, small GTPase signaling, and actin cytoskeleton organization (Fig. 4a), indicating extensive immune activation and transcriptional remodeling. Consistent with these transcriptional differences, histological analysis at 4 wpi revealed increased accumulation of spinal cord-associated T cells in recovered compared with non-recovered animals (Fig. 4e, f). The convergence of these transcriptional and cellular differences at 4 wpi, coinciding with pronounced differences in locomotor recovery and axonal regrowth, suggests that T-cell-mediated processes may contribute to the establishment of a regeneration-permissive environment.

**Figure 4.**
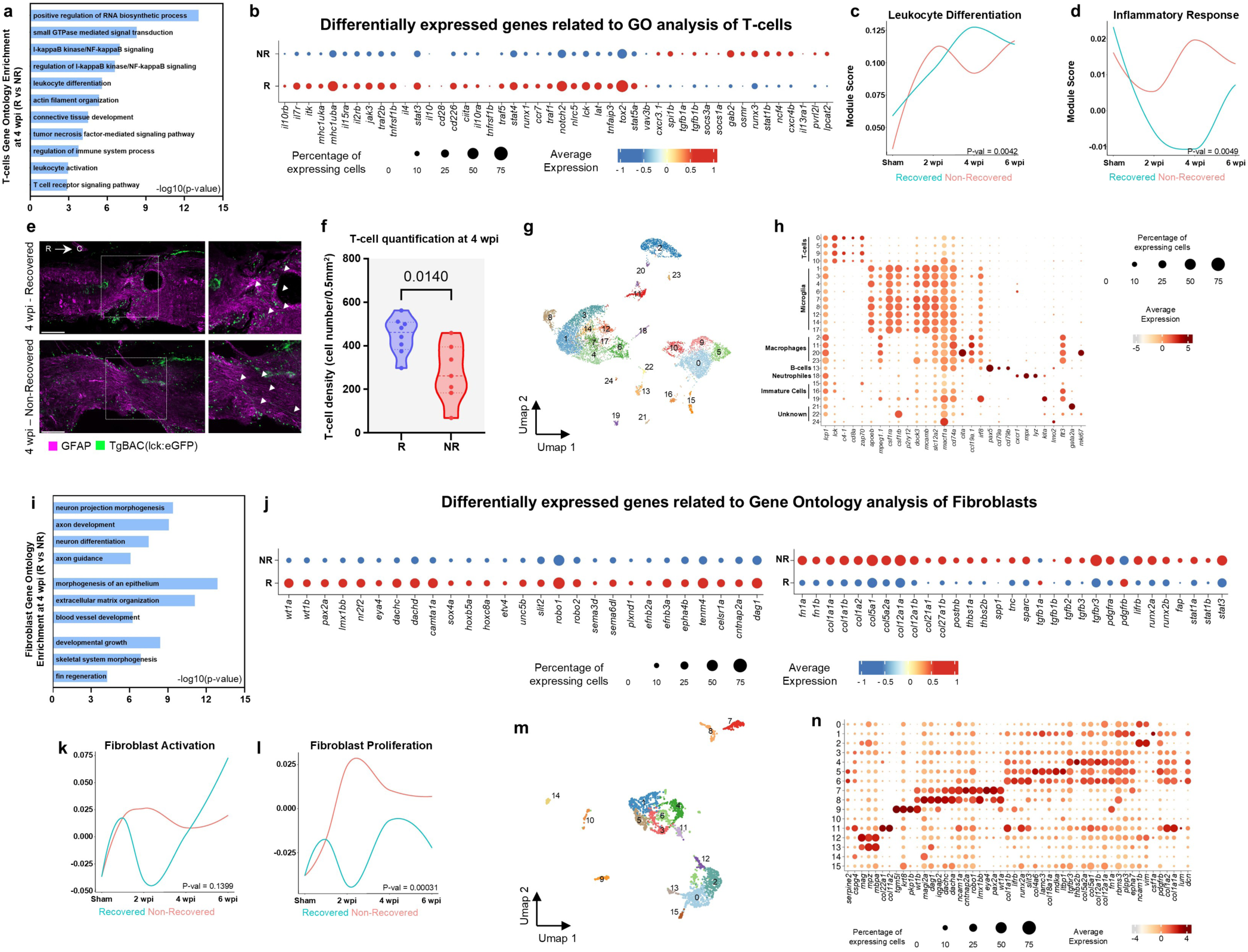
Successful spinal cord regeneration couples T-cell activation with developmental fibroblast remodeling. **A)** Gene Ontology Biological Process (GO-BP) enrichment analysis of differentially expressed genes (DEGs) in T cells from recovered (R) versus non-recovered (NR) animals at 4 weeks post-injury (wpi). Enriched terms included leukocyte differentiation, T-cell receptor signaling, regulation of immune system processes, NF-κB signaling, and positive regulation of RNA biosynthetic processes. **B)** Dot plot showing representative DEGs associated with the enriched GO-BP terms in T cells at 4 wpi. Dot size indicates the proportion of expressing cells and color represents scaled average expression. **C, D)** Temporal dynamics of T-cell transcriptional programs across the regeneration time course. Conditional means were smoothed using LOESS regression and plotted separately for R (blue) and NR (red) animals. Curves represent the activity of gene signatures associated with the indicated biological processes. Statistical significance was assessed by comparing the temporal trajectories between conditions. **E)** Representative longitudinal spinal cord sections from *lck:EGFP* animals stained with anti-GFAP antibody (magenta) at 4 wpi. T cells are shown in green. Arrowheads indicate representative T cells located within or adjacent to the lesion. Insets show higher magnification views of the boxed regions. Scale bars, 100 µm. **F)** Quantification of T-cell density within the injured spinal cord at 4 wpi. Recovered animals exhibited a greater density of T cells within the lesion environment compared with non-recovered animals (two-tailed unpaired *t*-test, p = 0.0140). **G)** UMAP representation of immune-cell subclustering revealing 25 transcriptionally distinct immune populations (clusters 0–24). **H)** Dot plot displaying marker gene expression used to assign immune-cell identities to the immune subclusters shown in **(G)**. Dot size indicates the fraction of expressing cells and color denotes scaled average expression. **I)** GO-BP enrichment analysis of DEGs in fibroblasts from R versus NR animals at 4 wpi. Enriched terms included neuron projection morphogenesis, axon development, axon guidance, developmental growth, extracellular matrix organization, blood vessel development, and regenerative developmental programs. **J)** Upper dot plot showing representative DEGs associated with developmental and regeneration-related transcriptional programs enriched in fibroblasts from R animals at 4 wpi. Lower dot plot showing representative extracellular matrix (ECM)-associated genes differentially expressed between R and NR fibroblasts at 4 wpi. Dot size indicates the fraction of expressing cells and color denotes scaled average expression. **K, L)** Temporal dynamics of fibroblast activation and fibroblast proliferation signatures across the regeneration time course. Conditional means were smoothed using LOESS regression and plotted separately for R (blue) and NR (red) animals. Statistical significance was determined by comparing temporal trajectories between conditions. **M)** UMAP representation of fibroblast subclustering identifies 16 transcriptionally distinct fibroblast populations (clusters 0–15). **N)** Dot plot displaying marker gene expression used to define fibroblast subcluster identities shown in **(M)**. Dot size indicates the proportion of expressing cells and color represents scaled average expression.

Integration of differential expression and GO analyses identified a distinct recovery-associated T-cell state emerging at 4 wpi (Fig. 4b-d). T cells from spinal cords of recovered animals showed elevated expression of genes involved in T-cell activation, signaling, differentiation and survival, including *traf1, traf5, vav3b, lat, itk, cd226, il21r, il7r, notch2,* and *bcl11ba*, together with genes associated with antigen presentation, including *mhc1uba, mhc1uka,* and *nlrc5* (Fig. 4b-d). In contrast, T cells from spinal cords of non-recovered animals preferentially expressed a limited set of genes associated with inflammatory signaling and immune stress responses, including *osmr, stat1b, gab2, spi1b, ncf4,* and *lpcat2*, and they failed to exhibit enrichment of coherent biological programs (^38–40)^.

To further resolve immune heterogeneity, immune cells were subclustered into 25 transcriptionally distinct populations comprising resident microglia, infiltrating macrophages, multiple T-cell states, B cells, neutrophils, and rare progenitor-like immune populations (Fig. 4g, h). Analysis of subcluster abundance revealed progressive remodeling of the immune compartment during regeneration. Although macrophage and microglial populations remained predominant throughout regeneration, recovered animals displayed increased representation of multiple T-cell populations beginning at 2 wpi (Fig. 2f) and persisting through 4 wpi (Fig. 2f, 4e). In contrast, spinal cord tissue of non-recovered animals remained enriched in myeloid populations and exhibited a late expansion of B-cell-associated and antigen-presenting populations at 6 wpi. Together, these transcriptional and compositional analyses indicate that successful regeneration is associated with coordinated activation and expansion of T-cell populations, whereas regenerative failure is characterized by persistent myeloid dominance and the absence of coordinated T-cell activation.

Fibroblasts also adopted distinct transcriptional programs according to regenerative outcome. At 4 wpi, GO analysis of fibroblasts from recovered animals showed enrichment for biological processes associated with axon development, axonogenesis, axon guidance, neuron projection morphogenesis, developmental growth, and regeneration, consistent with the acquisition of a sustained pro-regenerative transcriptional state (Fig. 4i). Correspondingly, these cells showed increased expression levels of multiple developmental regulators, including *wt1a, pax2a, lmx1bb,* and *tenm4* (^41,42^), whereas fibroblasts from non-recovered animals preferentially expressed extracellular matrix remodeling genes, including *fn1a, col12a1a, col12a1b, col27a1b, col5a2a, thbs1a, lifrb,* and *tnfaip2b* (Fig. 4j). By 6 wpi, these transcriptional differences were less prominent, suggesting that the critical divergence occurs during the intermediate phase of regeneration (Fig. 4k, l).

To further resolve stromal heterogeneity, fibroblasts were subclustered into 16 transcriptionally distinct populations (Fig. 4m, n). While all subclusters retained core extracellular matrix-associated features, several exhibited specialized transcriptional programs. In particular, fibroblast states enriched for developmental and axon guidance genes, including *wt1a, pax2a, eya4, lmx1bb,* and *robo1*, were preferentially expanded in recovered animals at 4 wpi, whereas non-recovered animals accumulated fibroblast populations enriched for structural extracellular matrix genes. Although these differences became less pronounced by 6 wpi, they indicate that regenerative outcome is associated with selective expansion of distinct fibroblast states rather than uniform activation of the fibroblast compartment.

Given the emergence of distinct immune and fibroblast states during regeneration, we next asked whether these transcriptional differences were accompanied by remodeling of intercellular communication using the CellChat pipeline (^43,44^). Overall, recovered and non-recovered spinal cords retained similar intercellular communication networks at both 2 and 4 wpi, with most signaling pathways detected in both conditions (Fig. S4). However, comparison of pathway activity revealed progressively greater differences over time, with substantially more pronounced pathway-specific rewiring at 4 wpi than at 2 wpi. These findings indicate that regenerative outcome is associated with selective remodeling of individual signaling pathways rather than large-scale changes in the overall communication network. Spinal cord cells in recovered animals preferentially engaged signaling pathways associated with neuronal growth, axon guidance, and cell adhesion, including NCAM, Netrin, Slit, Semaphorin, and Ephrin signaling, whereas non-recovered tissue exhibited relatively greater activity of pathways linked to extracellular matrix remodeling, inflammation, and tissue organization, including TGFβ, CXCL, IL16, FN1, and Thrombospondin signaling. Together, these findings indicate that successful regeneration is accompanied by coordinated remodeling of the signaling environment toward a pro-regenerative state, whereas regenerative failure is associated with the persistence of signaling programs consistent with chronic tissue remodeling and inflammation.

### Transcriptional regulators associated with divergent regenerative outcomes

Because differential transcriptional states ultimately arise through changes in transcriptional regulation, we next reconstructed cell type-specific gene regulatory networks (GRNs) using CellOracle (^45^) to determine how regulatory architecture differed between recovered and non-recovered spinal cords. As a prior framework for potential TF–target relationships, we used the prebuilt zebrafish promoter-based GRN provided with CellOracle, which is derived from promoter sequence and transcription factor-binding motif information. Thus, sample-specific chromatin accessibility data were not required for this analysis. CellOracle then integrated this base regulatory network with our scRNA-seq data to infer condition- and cell type-specific regulatory relationships at 2 and 4 wpi. Significant regulatory interactions were defined as predicted transcription factor (TF)–target edges with p < 0.05 and classified as shared between outcomes, recovered-specific, or non-recovered-specific. Overall GRN architecture remained broadly conserved between regenerative outcomes, with most predicted interactions retained in both conditions at 2 and 4 wpi (Fig. 5a; Table S4). Nevertheless, every cell population exhibited substantial condition-specific remodeling. Excluding the highly conserved neuronal network, approximately 25–35% of TF–target interactions were detected exclusively in one regenerative outcome (Table S4). Fibroblast GRNs were among the most extensively remodeled, with only 65.9% and 66.4% of interactions shared between recovered and non-recovered animals at 2 and 4 wpi, respectively, whereas T-cell networks became progressively rewired over time, decreasing from 74.5% shared interactions at 2 wpi to 67.9% at 4 wpi. In contrast, the consensus neuronal GRN remained highly conserved, retaining nearly 90% shared interactions at both time points (Table S4). These findings indicate that distinct regenerative outcomes arise through selective remodeling of otherwise conserved transcriptional networks, with regulatory rewiring preferentially affecting non-neuronal populations involved in shaping the injury microenvironment.

**Figure 5.**
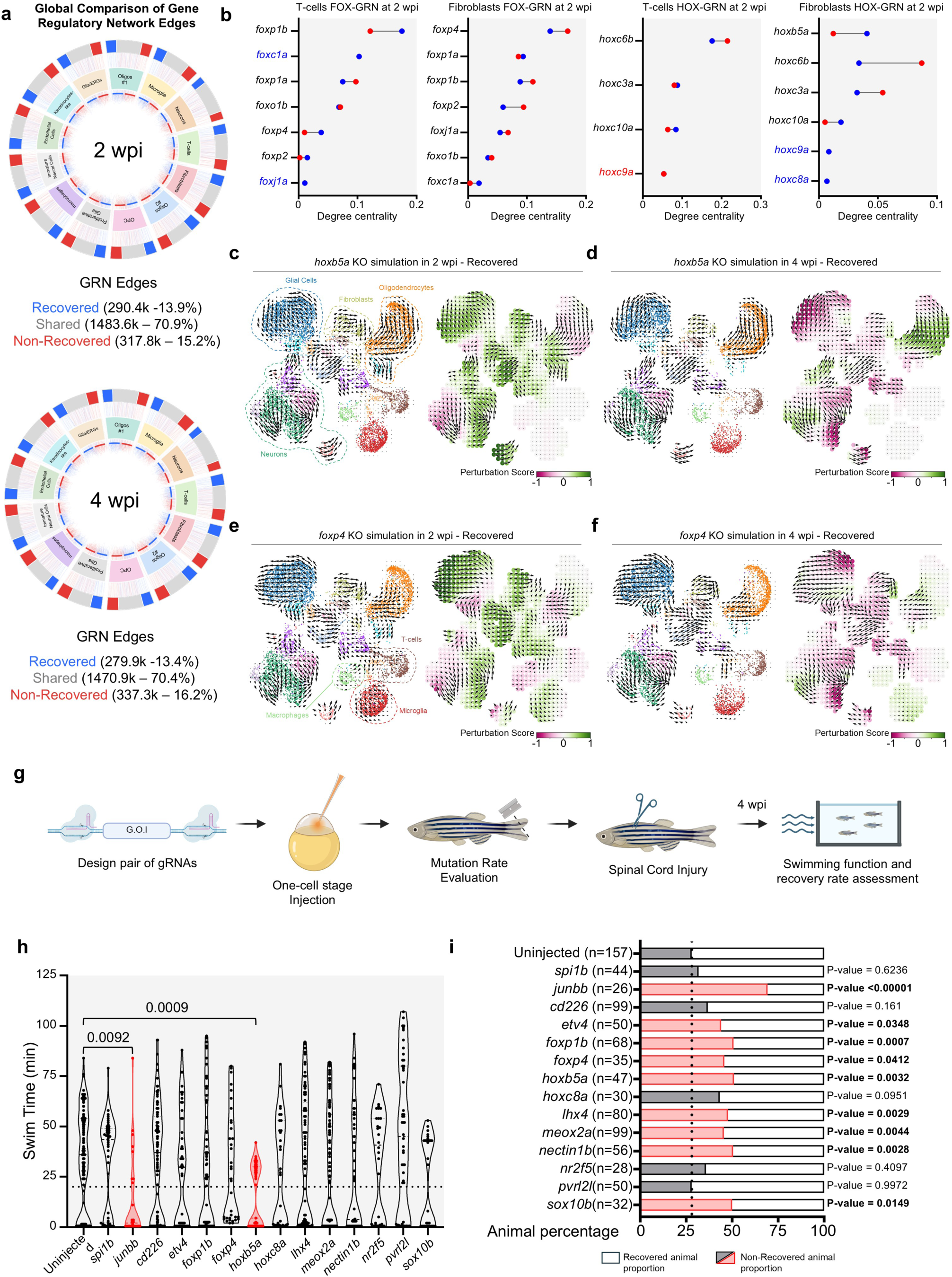
Regulatory network remodeling and functional screening reveal transcription factors controlling regeneration outcomes. **A)** Circos plots showing the conservation and condition-specific remodeling of CellOracle- inferred GRNs at 2 wpi and 4 wpi. Each sector represents a cell type, with inner arcs indicating recovered (blue) and non-recovered (red) animals. The outer ring summarizes GRN edges classified as recovered-specific (blue), shared between conditions (gray), or non-recovered- specific (red), with sector width proportional to the total number of significant TF–target interactions. Connecting bundles illustrate the distribution of regulatory interactions between each condition and the corresponding edge category, with bundle thickness proportional to the number of TF–target edges. Significant regulatory interactions were defined as CellOracle edges with p < 0.05. **B)** Cleveland plots comparing the degree centrality of expressed transcription factors of the FOX and HOX family between recovered (blue) and non-recovered (red) animals at the indicated time point. Within each panel, transcription factors are ranked according to their degree centrality in recovered animals, and paired values are connected by gray lines. Values shown correspond to CellOracle-derived degree centrality in T cells and fibroblasts. Δ values indicate the difference in degree centrality between recovered and non-recovered animals (recovered − non-recovered). Transcription factor highlighted in blue or red indicates expression limited to recovered (blue) or non-recovered (red) animals. **C-F)** Representative CellOracle knockout (KO) simulation of two transcription factors showing the highest degree centrality in fibroblasts at 2 wpi from the FOX and HOX gene family. **(C, D)** CellOracle KO simulation of *hoxb5a* at 2 **(C)** and 4 wpi **(D)**. **(E, F)** Cell Oracle KO simulation of *foxp4* at 2 **(E)** and 4 wpi **(F)**. For each KO simulation, data are shown as KO-induced, simulated cell state transition vector field projected on a UMAP highlighting cell-identity in dashed line (left panel), or simulated cell state transition vector field visualized with perturbation scores (PS; magenta: negative score; green: positive score) (right panel) **G)** Schematic representation of the experimental workflow for F_0_ CRISPR/Cas9-based functional screening of transcription factors identified through CellOracle KO simulation. For each target gene, a pair of gRNAs was designed for loss-of-function. gRNA pairs were injected into one-cell stage embryos that were subsequently raised until adulthood. Tail fin genotyping was performed to assess mutation presence in adult animals, then mutated animals underwent spinal cord transection and were tested for swim ability and recovery rate at 4 wpi. **H)** Swim capacity of F_0_ injected adult animals at 4 wpi. Data are shown as violin plot, each dot represents an individual fish. Dataset highlighted in red denotes significant differences from uninjected animals. Two-way repeated-measures ANOVA tests with Holm-Šidák correction were used; p-values less than 0.05 are shown for corresponding data. **I)** Stacked bar plots showing the percentage of recovered (white) and non-recovered (gray/red) animals in uninjected and F_0_ crispant animals. The total number of animals analyzed is indicated on the left side of each bar. Recovery status was determined based on swim tunnel endurance performance using the predefined recovery threshold described in Methods. Statistical differences in recovery frequency among genotypes at each time point were assessed using a χ² test of independence, followed by pairwise comparisons where appropriate. Exact p-values for gene crispants compared to uninjected populations are displayed on the right side of each bar. Statistically significant changes (p < 0.05) in non-recovered proportion are highlighted in red.

Although most regulatory interactions were preserved, network remodeling substantially altered the predicted influence of individual transcription factors. We therefore compared TF degree centrality between recovered and non-recovered fibroblast and T-cell networks (Fig. 5b; Table S4). Members of the FOX family emerged as recurrently rewired regulatory hubs and displayed the largest mean condition-dependent changes in degree centrality among the major TF families represented in these cell populations (Table S4). This effect was particularly evident at 4 wpi, when multiple FOX factors ranked among the most differentially connected regulators in both fibroblasts and T cells. Rather than exhibiting coordinated changes in a single direction, individual FOX factors displayed reciprocal alterations in network connectivity, consistent with redistribution of regulatory influence within the family. Several HOX and other homeobox transcription factors also underwent marked condition-dependent changes in centrality. In particular, *hoxb5a* represented the most highly connected canonical HOX factor in recovered fibroblasts at 2 wpi, whereas *foxp4* consistently ranked among the largest differences in network connectivity between regenerative outcomes (Fig. 5b; Table S4). Together, these analyses identified individual FOX and HOX transcription factors as previously unexplored candidate regulators associated with successful spinal cord regeneration.

To determine whether these changes reflected remodeling of downstream regulatory programs, we next compared the predicted regulons of *hoxb5a* and *foxp4* in fibroblasts. Despite retaining large conserved regulatory cores, both TFs displayed extensive condition-specific rewiring of their predicted target networks (Tables S4). At 2 wpi, the *hoxb5a* regulon comprised 1,485 interactions shared between recovered and non-recovered animals, together with 373 additional interactions detected only in recovered animals and 279 detected only in non-recovered animals. Similarly, the *foxp4* regulon contained 1,250 shared interactions, together with 254 recovered-specific and 346 non-recovered-specific interactions. Similar levels of rewiring persisted at 4 wpi, indicating that approximately one-third of the combined regulons of both transcription factors consisted of outcome-specific regulatory interactions (Tables S4). These findings suggest that differences in regenerative success arise not only from altered connectivity of individual transcription factors but also through extensive remodeling of the downstream transcriptional programs they regulate. To investigate whether these rewired regulatory hubs could influence regenerative cell-state dynamics, we performed CellOracle in silico loss-of-function simulations for *hoxb5a* and *foxp4* in recovered fibroblasts (Fig. 5c-f). Both perturbations produced stage-dependent effects on the predicted fibroblast transition landscape. At 2 wpi, knockout-induced transition vectors were predominantly aligned with the endogenous state-transition field and associated with positive perturbation scores, whereas at 4 wpi the predicted transition vectors opposed the endogenous trajectory and produced predominantly negative perturbation scores (Fig. 5c-f). These simulations suggest that the regulatory influence of *hoxb5a* and *foxp4* evolves throughout regeneration and depends on the underlying fibroblast state. Collectively, the GRN and perturbation analyses identified dynamically rewired transcriptional regulators with the potential to control regenerative fibroblast programs, providing a focused set of candidates for experimental tests.

### Hoxb5a is essential for spinal cord regeneration

To determine whether transcriptional regulators identified through GRN analysis were functionally required for successful spinal cord regeneration, we performed a targeted F_0_ CRISPR/Cas9 screen of candidate genes associated with regeneration-linked cellular states (Fig. 5g). Candidates were prioritized based on differential expression, GRN remodeling, predicted regulatory importance and known or predicted roles in developmental patterning, morphogenesis or immune regulation. For each target gene, two guide RNAs targeting promoter-proximal and coding regions were co-injected with Cas9 at the one-cell stage to generate predicted loss-of-function alleles. Adult crispants were genotyped from tail fin biopsies before undergoing complete spinal cord transection, and regenerative outcome was assessed at 4 wpi using the swim tunnel assay, allowing both continuous measurement of swimming endurance and classification of animals as recovered or non-recovered (Fig. 5h, i). Depending on mutagenesis efficiency and clutch availability, 16 to 99 animals were analyzed for each candidate.

Most candidate perturbations produced either no detectable phenotype or only modest effects on regeneration (Fig. 5h, i). Consistent with previous reports, disruption of *spi1b* did not significantly alter regenerative outcome compared with uninjected controls, supporting the specificity of the screening paradigm (^29^). In contrast, disruption of several developmental transcription factors increased the proportion of non-recovered animals, with *junbb* (implicated also in a previous crispant screen ^29^), *foxp1b*, and *hoxb5a* producing the strongest effects (Fig. 5h, i). Notably, *hoxb5a* emerged independently from multiple levels of our computational analyses, including differential expression, GRN centrality, regulon remodeling, and CellOracle perturbation analysis. Moreover, among GRN-derived candidates, *hoxb5a* crispants exhibited one of the strongest reductions in functional recovery, showing both a significant decrease in swimming endurance and an increased proportion of non-recovered animals compared with uninjected controls (Fig. 5h, i). Although several additional candidates including *foxp4* and *etv4* displayed trends toward impaired regeneration, these effects were less pronounced.

To reduce effects of genetic mosaicism, we generated stable mutant lines for 7 GRN-predicted candidate genes (*hoxb5a*, *etv4, hoxc8a, lhx4, nr2f5, cd226* and *pvrl2l*). While each of these mutants was viable to adulthood in the homozygous state, only *hoxb5a* mutants exhibited measurable impairments in swimming endurance following spinal cord injury (Fig. S5, Fig. S6a, b). While wild-type animals progressively recovered swimming endurance between 2 and 6 wpi, *hoxb5a* homozygous mutants displayed a persistent reduction in swimming performance throughout regeneration timepoints (Fig. 6a, b). These functional deficits translated into a markedly higher proportion of non-recovered animals, with ∼76% versus ∼37% at 2 wpi, ∼59% versus ∼18% at 4 wpi, and ∼63% versus ∼13% at 6 wpi for *hoxb5a* mutants and wild-type animals, respectively (Fig. 6c). Together, these findings establish that Hoxb5a is a significant factor in promoting efficient functional recovery after spinal cord injury.

**Figure 6.**
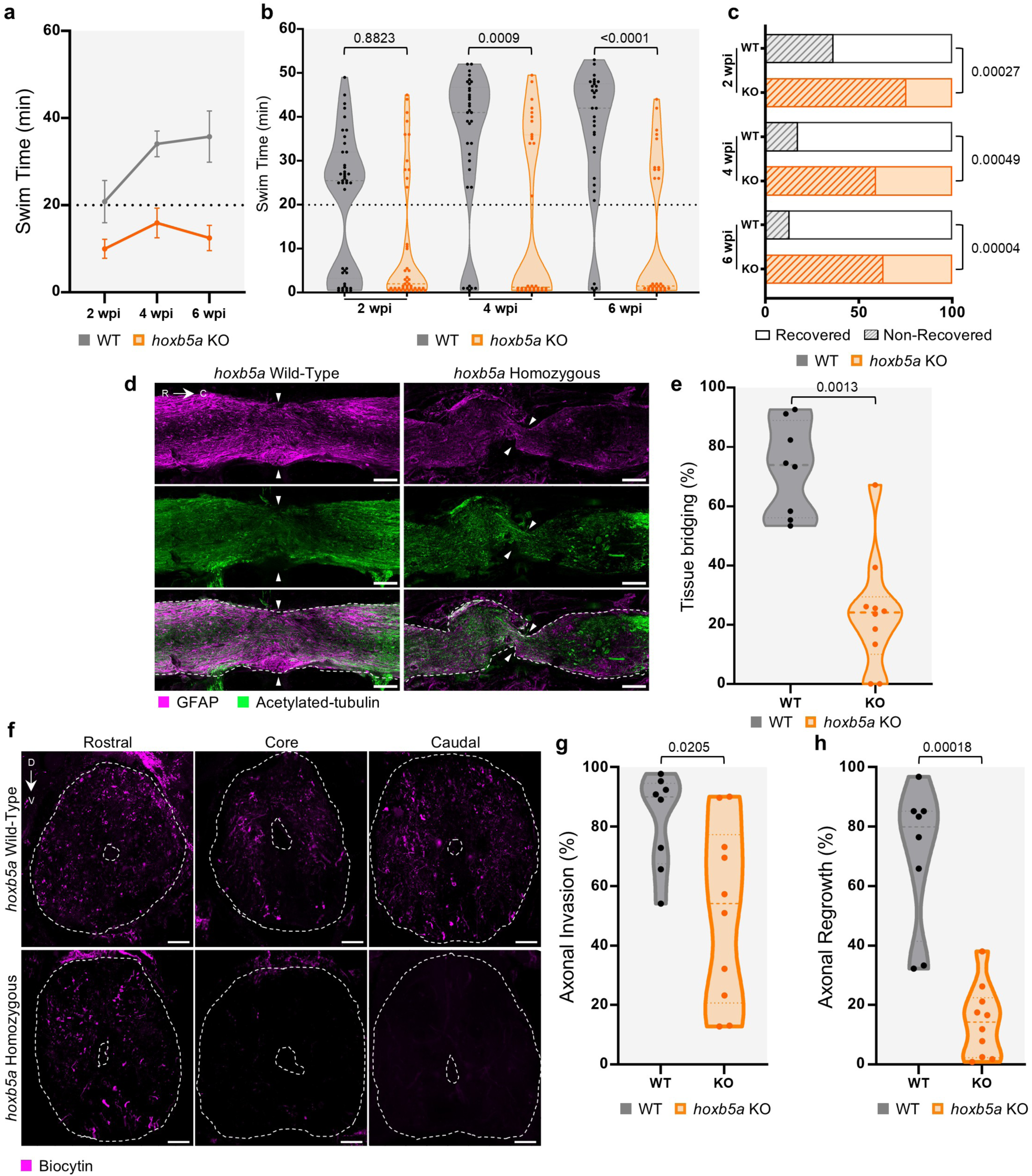
Loss of *hoxb5a* impairs tissue bridging, axon crossing and functional recovery after spinal cord injury. **A, B)** Swim capacity of wild-type (WT - gray) and homozygous *hoxb5a* mutants (KO - orange) at increasing current over a 6-week follow-up period after spinal cord transection. Data are shown as line graph **(A)** at 2, 4 and 6 wpi and violin plot **(B)** at the similar timepoint. Each dot represents an individual fish. Dashed line indicates the 20 min acclimation period and serves as a threshold to identify non-recovered animals. Error bars in **(A)** indicate SEM. Two-way repeated-measures ANOVA tests with Holm-Šidák correction were used to calculate exact p-values, only significant p-values (p < 0.05) are shown for corresponding data. **C)** Proportion of recovered and non-recovered animals following spinal cord injury. Stacked bar plots showing the percentage of recovered (clear bar) and non-recovered (dashed filled bar) animals in wild-type (WT - white) and *hoxb5a* knockout (KO - orange) groups at 2, 4, and 6 wpi. Statistical differences in recovery frequency among genotypes at each time point were assessed using a χ² test of independence. Exact p-values from comparisons with WT are displayed on the right of corresponding bar. Sample sizes were: 2 wpi, WT (n = 41), KO (n = 45); 4 wpi, WT (n = 34), KO (n = 32); 6 wpi, WT (n = 31), KO (n = 30). **D, E)** Analysis of tissue bridging following spinal cord transection in *hoxb5a* mutants. Representative longitudinal tissue sections indicating the glial marker GFAP (Magenta) and axonal marker acetylated α-tubulin (green) in WT and *hoxb5a* KO at 4 wpi are shown. n = 10-15. Quantification of tissue bridge thickness is shown on the right side **(E)**. A two-tailed Mann– Whitney test was used for comparisons. Scale bars, 100 µm. **F-H)** Analysis of axonal invasion and regrowth after anterograde axon tracing at 4 wpi. Representative transverse sections rostral, at the core of, and caudal to the injury site are shown, with biocytin-labelled axons (magenta) and a white dashed line highlighting spinal cord and central canal boundaries. Quantification is to the right. A two-tailed Mann–Whitney test was used for comparisons. n = 8-10. Scale bars, 50 µm.

To determine whether impaired recovery was associated with altered tissue repair, we next examined tissue bridge formation at 4 wpi by immunostaining longitudinal spinal cord sections for glial and axonal tissue (Fig. 6d,e). Homozygous *hoxb5a* mutants exhibited a profound reduction in tissue bridge formation, with regenerated tissue spanning ∼24% of the spinal cord width compared with ∼73% in wild-type animals (Fig. 6e). To quantify axonal regeneration in wild-type and knockout animals, we used anterograde biocytin tracing at 4 wpi (Fig. 6f-h; Fig. S6c). Wild-type animals displayed robust axonal invasion into the injury core together with extensive regrowth beyond the lesion. In contrast, *hoxb5a* knockout animals exhibited markedly reduced axonal invasion within the lesion, with an invasion index of ∼51% compared with ∼82% in wild-type animals, together with severely impaired axonal regrowth beyond the injury site (∼14% versus ∼70%, respectively), indicating that loss of *hoxb5a* profoundly compromises successful axon crossing following spinal cord injury. Thus, empirical tests of GRN regulators determined by comparison of regenerative failure and success contexts reveal an important role for *hoxb5a* in axon pathfinding and functional recovery during innate spinal cord repair.

## Discussion

Differences in regenerative capacity have been extensively documented between species with widely different regenerative capacity, as well as across developmental stages and aging (^41,46,47^). Here, we analyze a substantial and reproducible subset of adult zebrafish that spontaneously fail to recover locomotor function following identical spinal cord injuries. Importantly, this divergence establishes a single-species model for identifying endogenous mechanisms that determine regenerative competence, avoiding potential issues with cross-species comparative models.

Our findings suggest that failure of axon crossing in non-recovered animals reflects a qualitative defect in the lesion environment rather than an inability to restore tissue continuity. Fibroblasts in these animals retain extracellular matrix remodeling programs reminiscent of the fibrotic environment that restricts axonal growth after mammalian spinal cord injury (^5,48^), whereas those in recovered animals preferentially express developmental guidance molecules, including *robo1/2, slit2, sema3d, sema6dl* and *epha4b*. The cellular enrichment of *epha4b* and *sema6dl* in recovered animals likely spatially organizes these guidance cues, given that manipulation of EphA4 and Sema6D signaling in mammalian spinal cord injury alters axonal retraction and regenerative growth (^49,50^). Axon growth across the lesion, however, is only the first step toward functional recovery, as studies suggest that boosting growth across an injury is insufficient to restore locomotion unless regenerating neurons are directed toward relevant downstream targets by strategic guidance cue placement (^51,52^). Our findings highlight the importance of defining a minimal pattern of reconnection required for recovery, particularly as functional circuits may be rebuilt through new relay pathways rather than exact restoration of their original architecture (^53–56)^. More broadly, anatomical repair does not necessarily predict functional restoration: peripheral nerves can regenerate across a repaired lesion yet remain functionally impaired because of axonal misrouting (^57^), while healed skin and tendon can regain tissue continuity without fully restoring native structure or biomechanical function (^58,59^). Thus, our results emphasize that successful regeneration requires not only structural repair, but also reconstruction of a molecular environment and connectivity pattern capable of restoring function.

Gene regulatory network analysis enabled comparison of regulatory organization between two closely related injury responses that ultimately produce opposite regenerative outcomes. Similar network-based approaches have distinguished successful from failed cell-state transitions and identified regulatory determinants that were not apparent from differential expression alone (^45,60^), supporting their use as a framework to prioritize candidate regulators for functional testing. These predictions remain hypothesis-generating, and gene perturbations, ideally toward full germline mutants, are critical. In our study, this combined approach revealed Hoxb5a as important for spinal cord regeneration, yet single gene perturbations may underestimate functional redundancy or cooperation among transcription factors. Such redundancy is well established within the HOX family: in recent studies, the full patterning functions of the Hox10 and Hox11 paralogous groups became apparent only after simultaneous disruption of multiple family members, whereas combined loss of *Hoxa11* and *Hoxd11* produces substantially more severe limb-patterning defects than either single mutation (^61,62^). Similar combinatorial control may operate during regeneration, particularly among developmental patterning factors identified by our network analyses, which could explain why disruption of several individual candidates did not produce gross phenotypes. Multiplexed perturbation approaches have already demonstrated the value of resolving genetic interactions at single-cell resolution (^63,64^), and adapting such strategies to regeneration could help define the minimal regulatory combinations required to control regenerative outcome.

The involvement of *hoxb5a* in spinal cord regeneration is intriguing given the broader relationship between HOX genes and positional information. Adult fibroblasts retain aspects of the embryonic HOX code that reflect their anatomical origin (^65–68)^, and this positional information can remain functionally instructive: depletion of *HOXA13* in human distal fibroblasts disrupts their ability to impose site-specific epidermal identity (^67,68^). Importantly, HOX activity in mesenchymal cells may have additional or alternative roles. Hox genes canonically act through overlapping, context-dependent expression domains, and Hox5 paralogs can regulate cell positioning, matrix adhesion and extracellular matrix organization without overtly altering fibroblast identity (^69^). In zebrafish, *hoxb5a* is expressed in multiple embryonic mesodermal compartments and contributes with other *hoxb* genes to positional patterning of the lateral plate mesoderm (^70^), raising the possibility that a similar developmental program is redeployed during axial structure regeneration. This possibility is particularly relevant in fibroblasts, where *hoxb5a* emerged as a regulatory hub and where some of the strongest differences between successful and failed regeneration were observed. Fibroblasts of non-recovered animals retained ECM remodeling programs, whereas those from recovered animals preferentially expressed developmental and axon growth-associated programs. Consistent with a role in organizing this stromal response, *hoxb5a* mutants showed reduced tissue bridge formation together with impaired axonal invasion and regrowth. Thus, one mechanism by which *hoxb5a* might facilitate axonal regeneration is by regulating the fibroblast ECM and signaling environment through which regenerating axons extend.

## Materials and methods

### Zebrafish lines and husbandry

Zebrafish strains of the Ekkwill (EK) genetic background at the age of 4 to 6 months were used for this study. Animals were housed at 28°C with a light/dark cycle of 14:10 hours at a density of 15 to 18 animals per 3L tanks. The following published transgenic line was used for this study: TgBAC(lck:eGFP) (^71^). Experimental procedures performed on adult zebrafish were approved by the Institutional Animal Care and Use Committee (IACUC) at Morgridge Institute for Research and UW-Madison. Control and experimental animals were maintained under identical housing, feeding and husbandry conditions throughout the study. Adult male and female zebrafish were used in approximately equal proportions.

### Spinal cord transection

Spinal cord transections on adult zebrafish were performed as previously reported (^16^). Briefly, zebrafish were anesthetized using 0.02% aminobenzoic-acid-ethyl methyl-ester (MS222, Sigma) in fish water. Fine scissors were used to make a small incision through skin and muscle and expose the vertebral column. Transection injuries were performed halfway between the brainstem and the caudal fin. Complete transection was confirmed visually by complete separation of the spinal cord stumps and absence of residual tissue connecting the lesion margins. Zebrafish were allowed to recover at 28 °C. Sham-injured fish underwent an incision through skin and muscle to expose the spinal cord, which was left intact.

For biocytin treatment used for anterograde axon tracing, adult fish were anaesthetized using 0.02% aminobenzoic-acid-ethyl methyl-ester (MS222) in fish water. A secondary spinal cord transection was performed 2 mm rostral to the initial spinal cord injury. A biocytin-soaked gelfoam surgical sponge was inserted at the new injury site (Gelfoam, Pfizer, 09-0315-08; Alexa Fluor 594 Biocytin, Thermo Fisher Scientific A12922). The incision was closed and glued using Vetbond tissue adhesive material (Santa Cruz Biotechnology, sc-361931). Tissue was collected 24 hrs after biocytin application. Biocytin-positive axons were quantified in Fiji (ImageJ) using the Threshold and Analyze Particles functions under identical thresholding parameters for all images. Particle counts from the two rostral sections were averaged and used as an internal normalization control to account for differences in biocytin uptake and tracing efficiency between animals. The axon invasion index was calculated as the ratio of the average number of biocytin-positive particles within the lesion core to the average particle count in the rostral control sections. Similarly, the axon regrowth index was calculated as the ratio of the average number of biocytin-positive particles in the caudal spinal cord to the average particle count in the rostral control sections. All image acquisition and quantification were performed blinded to the experimental group. All images were acquired using a Zeiss LSM 900 confocal microscope.

### Behavioral assays

Swim capacity recovery after spinal cord transection was measured by exercising fish in groups of 8 to 12 animals in a 5L swim tunnel respirometer device (Loligo, SW100605L, 120 V/60 Hz). Fish were acclimated for 20 min at a fixed low water current in the enclosed tunnel (7cm/s), then water current velocity was increased every 2 min and fish swam against the current until they reached exhaustion. For each animal reaching exhaustion, swim time and current velocity at exhaustion were recorded for downstream analysis. Swimming capacity recovery assay was performed at 2, 4 and 6 weeks post-injury. Animals were classified as non-recovered (NR) if they were unable to maintain their position against the 7 cm s⁻¹ water current during the 20-min acclimation period of the swim tunnel assay. Animals were tested 3 days post-injury in groups of 5 to 8. Tested animals were subjected to a 10 min period at fixed low flow (7cm/s) and any animals displaying swimming ability at 3 days post-injury were excluded from follow-up experimental procedure and/or analysis. This category of animals represented approximatively 1% of performed injuries.

Swim ability in an undisturbed environment was quantified using a ZebraCube from Viewpoint Behavioral Technology. Eight individual animals were assessed simultaneously. Each animal was kept in an individual 2L tank positioned within the ZebraCube in the dark for a 15 min acclimation period before being recorded for 30 min. Swimming behavior tracking, including speed and distance parameters, were measured using the built-in software provided by ViewPoint Behavioral Technology (ViewPoint app manager – Tracking locomotion/gravity center version 5.33.0.250).

### Histological analysis

Dissected tissues were collected and fixed overnight at 4°C in a 4% PFA solution, then rinsed and washed in phosphate buffer solution with 0.5% Tween (Sigma). Samples were incubated in 30% sucrose solution at 4°C overnight for cryoprotection. Samples were then embedded in Tissue Freezing Medium (TFM – General Data Healthcare) and frozen on dry ice before being stored at -80°C until used.

Transverse cryosections (20–25 µm) were used for immunohistochemistry and axon tracing analyses, whereas 50-µm longitudinal cryosections were used for tissue bridge quantification. Tissue sections were incubated in 100% methanol at -20°C for 10 min; sections were then brought to room temperature, dried and rinsed using PBS-Tween. Sections were blocked and permeabilized in blocking buffer (0.2% Triton X-100, 0.5% Tween, 5% goat serum in PBS) for 4 h at room temperature. Sections were then incubated overnight at 4°C with primary antibodies diluted in blocking buffer. Primary antibodies were washed using PBS-Tween and incubated in secondary antibodies solution for 4 h at room temperature. Slides were washed using PBS-Tween at room temperature and then incubated with DAPI (ThermoFisher Scientific, D3571, 1:5000) at room temperature for 15 minutes. Following washes, sections were mounted using glycergel (Dako) and kept at 4°C until imaging. Primary antibodies used were: chicken anti-GFP (Aves Labs, GFP-1020, 1:500), mouse anti-GFAP (ZIRC, Zrf1, 1:200), rabbit anti-GFAP (Sigma, G9269, 1:200) and mouse anti-acetylated-α-tubulin (Sigma, T6793, 1:1000). Secondary antibodies used in this study were: Alexa Fluor 488/633/594 (1:200) goat anti-rabbit, anti-mouse or anti-chicken antibodies (Alexa Fluor 488 Goat anti-Rabbit, Thermo Fisher Scientific, A-11034; Alexa Fluor 594 Goat anti-Rabbit, Thermo Fisher Scientific, A-11037; Alexa Fluor 488 Goat anti-Mouse, Thermo Fisher Scientific, A-11029; Alexa Fluor 633 Goat anti-Mouse, Thermo Fisher Scientific, A-21050; Alexa Fluor 594 Goat anti-Mouse, Thermo Fisher Scientific, A-11032; Alexa Fluor 488 Goat anti-Chicken, Thermo Fisher Scientific, A-11039; Alexa Fluor 633 Goat anti-Mouse, Thermo Scientific, A-21052).

Tissue bridge formation was quantified on 50-µm longitudinal cryosections immunostained for GFAP and acetylated-tubulin. Images were acquired using identical confocal microscope settings for all samples. For each animal, three to five longitudinal sections containing the lesion center were analyzed. The tissue bridge was identified as the continuous GFAP-positive structure spanning the injury gap. Bridge thickness was measured at the midpoint of the lesion using Fiji (ImageJ) by drawing a line perpendicular to the longitudinal axis of the spinal cord across the narrowest region of the GFAP-positive bridge. Measurements from all analyzed sections were averaged to obtain a single value for each animal. Quantification was performed blinded to the experimental group. All images were acquired using a Zeiss LSM 900 confocal microscope.

### Spinal cord nuclei isolation

Adult zebrafish spinal cord tissue spanning approximately 2 mm rostral and caudal to the lesion center was dissected from sham-operated, recovered, and non-recovered animals at the indicated time points following spinal cord injury. For each biological replicate, spinal cords from 15 animals were pooled prior to nuclei isolation. Dissected spinal cords were flash frozen in liquid nitrogen and maintained at -80°C until used. Spinal cord nuclei were isolated following a previously published protocol (^72^). Briefly, frozen tissues were transferred to a pre-cooled Dounce homogenizer (Sigma) with 1 ml of 1x Homogenization Buffer (supplemented with 0.2 U/µl SUPERase RNAse inhibitor– ThermoFischer Scientific - and 1:200 dilution of PIC III – Sigma -) as described by Corces R. and colleagues (^73^). Samples were homogenized 10 times using an A Pellet Pestle and 6 times using a B Pellet Pestle and incubated 6 min on ice. Samples were then filtered through a 40 µm cell strainer and centrifuged at 300 g for 10 min at 4°C. Pellets were resuspended in PBS (supplemented with 1% BSA and 1U/µl of RNAse inhibitor). Resuspended nuclei were layered on top of a sucrose cushion (homogenization buffer with 2M sucrose) and centrifuged at 16,000 g for 30 min at 4°C to separate nuclei from cellular debris. Cleaned nuclei were then resuspended in PBS (supplemented with 2% BSA and 1 U/µl of SUPERase RNAse inhibitor). Nuclei concentration was determined by counting nuclei on a hemocytometer. Single nuclei profiling was performed using the Parse Biosciences Evercode Whole Transcriptomic kit. Nuclei fixation, barcoding, cDNA amplification and libraries were prepared following manufacturer instructions and sequenced on a DNBseq platform from BGI- Biosciences/Innomics to obtain 150 bp paired reads. Raw reads were mapped on the *Danio rerio* reference genome GRCz11. FASTQ files were processed using Trailmaker pipeline module (https://app.trailmaker.parsebiosciences.com/, pipeline v1.5.0, Parse Biosciences).

### Single nucleus sequencing analysis

Downstream analyses were performed in R (v4.4.3) using Seurat (v5.3.0). Genes detected in fewer than (20) nuclei were excluded from downstream analyses. Nuclei with fewer than (300) detected genes, greater than (3000) detected genes, or greater than 5% mitochondrial transcript content were removed during quality-control filtering. Putative doublets were identified using (doubletFinder) (v3) and excluded prior to downstream analyses. Filtered count matrices were normalized using the LogNormalize workflow implemented in Seurat. Highly variable genes were identified using the FindVariableFeatures function, and principal component analysis (PCA) was performed on the scaled expression matrix. Shared nearest-neighbor graphs were generated using the first 15 principal components, followed by unsupervised clustering using the Louvain algorithm at a resolution of 0.75. Uniform Manifold Approximation and Projection (UMAP) was used for dimensionality reduction and visualization. Cell-type identities were assigned based on canonical marker gene expression, differential gene expression analysis, and previously described spinal cord cell populations. Cluster marker genes were identified using the FindMarkers function in Seurat with default Wilcoxon rank-sum testing and Bonferroni-adjusted significance thresholds.

### Differential expression and pseudobulk analysis

To minimize cell-level pseudoreplication and account for biological replicate structure, differential expression analyses were performed using a pseudobulk strategy. Raw counts were aggregated by biological replicate, experimental condition, time point, and cell type using the “AggregateExpression” function in Seurat.

Differential expression analyses were performed independently for each cell population across regeneration conditions and time points using the Seurat FindMarkers function, including recovered versus non-recovered comparisons at 2, 4, and 6 weeks post injury, as well as comparisons to sham-operated controls. Genes with a min.pct >0.25, adjusted p-value < 0.05, and absolute log2 fold-change > 1 were considered significantly differentially expressed. Multiple testing correction was performed using the Benjamini–Hochberg procedure.

### Gene ontology and pathway enrichment analysis

Gene ontology (GO) enrichment analyses were performed on significantly differentially expressed genes using the Biological Process ontology. Over-representation analyses were conducted using clusterProfiler (v 4.14.4) with Benjamini–Hochberg multiple testing correction. GO terms with adjusted p-value < 0.05 were considered significantly enriched. Pathway enrichment analyses were additionally performed using KEGG annotations through clusterProfiler. Enrichment analyses were performed separately for upregulated and downregulated gene sets using all genes detected within each analyzed cell population as the background gene universe.

### Gene regulatory network inference using CellOracle

Gene regulatory network (GRN) analyses were performed using CellOracle (v 0.20.0) in Python (v 3.10.20). Processed Seurat objects were converted to AnnData format for downstream analyses. A prior zebrafish transcription factor regulatory network was constructed using promoter DNA sequence information and transcription factor motif databases implemented within CellOracle using the GRCz11 reference genome (^45^).

CellOracle was used to infer cell-state-specific GRNs by integrating transcription factor binding motif predictions with observed gene co-expression patterns across single nuclei. Separate GRNs were generated for recovered and non-recovered conditions at each post-injury time point. Network topology metrics, including degree centrality and betweenness centrality, were calculated for each transcription factor within individual cellular populations. To characterize regenerative-state-specific network rewiring, transcription factor network properties were compared between recovered and non-recovered conditions across time. Dimensionality reduction of GRN topology metrics was performed using principal component analysis (PCA) to visualize global network divergence between regenerative states. Candidate transcription factors were prioritized based on regenerative-state-specific changes in network centrality.

### CellChat analysis

CellChat 2.1.2 (^43,44^) was used on single nucleus data to measure cell to cell communication. The CellChat database is based on mouse or human genes; therefore, zebrafish genes were converted to mouse homologs. First the homolog list was obtained from Ensembl (https://www.ensembl.org/). Next zebrafish genes were mapped to their mouse homologs. For any mouse homolog that was duplicated among different individual zebrafish genes, the processed expression data for these genes were averaged to have one value for the mouse homolog, otherwise the data was directly transferred to the mouse homolog. If there is more than one mouse homolog for a zebrafish gene, the zebrafish expression value is applied to all the mouse homologs. The resulting expression matrix was turned back into a Seurat object, the meta data from previous annotation was added back in and then used for CellChat analysis and run separately on all time points for both recovered and non-recovered samples to calculate communication probabilities and likely ligand-receptor pairs and/or pathways that are communicating for a given cell type. Interactions were evaluated between conditions by merging the recovered and non-recovered for each time point. We ran “compareInteractions” to obtain overall weights of cell type to cell type interactions, “rankSimilarity” to find pathway distances, and “rankNet” to get the comparison of communication probabilities of pathways.

### Crispant and mutant line generation

For each gene targeted in the present study, 2 guide RNAs were designed using the CHOPCHOP website (^74^) aiming to generate large deletion. gRNAs were co-injected with 300ng/µl of Cas9 protein (PNA-Bio) into one-cell stage embryos. The efficiency of gRNAs was tested by genotyping 3-day post-injection embryos, and only gRNAs displaying a 90% mutation rate were used to generate crispants alleles used in this study. gRNA sequences and associated genotyping primers for each gene are listed below (5’ – 3’ orientation):

*<u>cd226</u>* (gRNA1: CACATAATCCGAGAAAAGAG**AGG**, Fw1: GTGCCTGATAAAAGTCCCCTAA, Rv1: TCCCTTTGAGAACAATCAATCA, gRNA2: TCCTCAGGGTTCATGGACCA**AGG**, Fw2: GAGGGAAGAGTGGAGTTTCTGA, Rv2: TTCGGGTTGAAATTTCCCTTTA),

*<u>etv4</u>* (gRNA1: ATCGCCGCTTGAGAATCCCA**AGG**, Fw1: CAAACGCTCAAAACAACCATAA, Rv1: ATGAAACCTCTCTGCTCTTTGC, gRNA2: GGACGCAGAATTACCCCCTC**AGG**, Fw2: TTTGTAGAGGTCGCAAGGAAAT, Rv2: GCTGAACAGTCCACATAAAAAGA),

*<u>foxp1b</u>* (gRNA1: CTGGCGACTCTAGTATTGCC**TGG**, Fw1: TGCATAAGGAAGCGTTCAGTAA, Rv1: TCGCGTGAACCAGTTGTATATC, gRNA2: GCCGCAAAAGATCAGCGGGT**AGG**, Fw2: GTGCGGCATAACCTTAGTCTTC, Rv2: TAACTCACATACGGGCACAAAG),

*<u>foxp4</u>* (gRNA1: CAACCCCCATCACCCCGATG**AGG**, Fw1: CTCTAGCTAAACCTGGCCTCAA, Rv1: CATGATGAGCTGTATTTGGTGG, gRNA2: CACAGCAACGTGGAAGGTAA**TGG**, Fw2: CACCTGCTCTCTTCCATTCTCT, Rv2: TCCTTGCTGAATGAAAGTGCTA),

*<u>hoxb5a</u>* (gRNA1: AACGCTTCGTACAGGGACTC**CGG**, Fw1: CCAAGATCTACAAATCAAGCCC, Rv1: GTTGAACGATTGACGCTTAGGT, gRNA2: TGGTTGGAAAGCGCTACCTG**CGG**, Fw2: CCCGAAGAAGGAGGATAGAGAT, Rv2: TAAAACATCCATAACATGCCGA),

*<u>hoxc8a</u>* (gRNA1: TACTGGTCATTTCAGTCCTG**CGG**, Fw1: TGTAGTCATTGTGGGTTATGGC, Rv1: TTTAATGGCCGCTTTAGAACAT, gRNA2: AATAGCACTAACCCTGGAGA**AGG**, Fw2: CTTTATGGTACCCAGCAAGAGG, Rv2: AGGCTACCCCACCTGTTTTAAT),

*<u>junbb</u>* (gRNA1: GGGTTACGGTCACAACGACG**CGG**, Fw1: AACCACCGGTTTTCAAACTCTA, Rv1: CCTGATAGAGGTCCGATTTGAG, gRNA2: GCAAACTAGAACGCATCGCG**CGG**, Fw2: AAAGAGGAACCGCAGACTGTT, Rv2: CTCAGGACCTTCTGCTTGAGTT),

*<u>lhx4</u>* (gRNA1: TGAACTCCGCTCTCTCTAGA**GGG**, Fw1: AGACTGAAGCCCGTGTTATTGT, Rv1: TGCTTAATGGCAGAAAAGGTCT, gRNA2: GGAGTGCCACTGCAACGTAA**GGG**, Fw2: GACTGGATGAAAATGATGCAAA, Rv2: ACAATTCCCCAGTGGAGTTTAAT),

*<u>meox2a</u>* (gRNA1: GCGGGGTGCACGGCCTGTGG**AGG**, Fw1: ATGCATACGGACACACTTCTTG, Rv1: AGAGATCCGGGTATGAGGAGAG, gRNA2: TCCCTGGAGGAGATTTCGGACGG**CGG**, Fw2: ATCATCATCATCATCATTTGGC, Rv2: AGAGCTGTCGCTTCTCCTTTTA),

*<u>nectin1b</u>* (gRNA1: GTCCGTTTTAGGTCACTAGT**AGG**, Fw1: AGGCTGTAACAAGAGGGAGATG, Rv1: TCCTCATATTTTCCCCTTCAAA, gRNA2: ATTGCGAGCGAACGGATGGT**CGG**, Fw2: ACTCTCAGCATTTCTCCGTTTC, Rv2: ATTGCATCTTTACGCAGGATTT),

*<u>nr2f5</u>* (gRNA1: AATAGTTCACGTTTTTGCCC**TGG**, Fw1: AGCAACGCCATTTTTATTTTGT, Rv1: TCTCAAGCAGAGACAAGCTCAG, gRNA2: AACCTGAGTTACACCTGCCG**GGG**, Fw2: AGACAAAATTCCCAATGTGGAC, Rv2: TACCCTCTCTTCTCATTCCGAC),

*<u>pvrl2l</u>* (gRNA1: TACGGTCCTCTTCAAGTGCT**CGG**, Fw1: TCACGAGCACAATAACATACCTG, Rv1: ATTACCTTGGCTGGACAGAAGA, gRNA2: GAAGACACGCGGCCAGAACC**AGG**, Fw2: CCTCGCAAAGTGAAAGAGATTT, Rv2: TGTTGTGAGGTAAACATCCTGC),

*<u>sox10b</u>* (gRNA1: GCCGAAATCACCGCCCTCTG**GGG**, Fw1: AAAGTAAGATGCCCCTGTGAGA, Rv1: TACTTTACAGTGCAGCAGGTCG, gRNA2: GGTCTCAGTGAAGTCCGACG**AGG**, Fw2: GGTCAGCAGTCTCAGATGTCC, Rv2: GATATTGATCCGCCAGTTTCCT),

*<u>spi1b</u>* (gRNA1: CAAGAGTTAAGCAAACAGCC**TGG**, Fw1: CATTTGAAAGGGAAATGATGGT, Rv1: GAGGCTGGTTTTGGAATTTATG, gRNA2: GAGGGATGTGATGGCTACCC**AGG**, Fw2: TTTACAGAGCTACAAAGCGTGC, Rv2: CGTCACAATGCACCAAATAAAC).

Stable mutant lines were generated for *hoxb5a, hoxc8a, etv4, nr2f5, lhx4, cd226* and *pvrl2l* using gRNAs listed above. The allele designations for these lines are: *pm1, pm2, pm3, pm4, pm5, pm6* and *pm7* respectively.

### Xenium spatial transcriptomics

For Xenium spatial transcriptomic analysis, spinal cord tissues were collected at 1, 2, 4 and 6 wpi, with recovered and non-recovered animals analyzed separately at 2, 4 and 6 wpi. Samples were fixed overnight at 4 °C in 4% paraformaldehyde (PFA; pH 7.3), washed five times for 15 min in PBS at room temperature, and cryoprotected for 4 h in RNase-free 30% sucrose. Tissues were then embedded in Tissue Freezing Medium, frozen, and stored at −80 °C until processing. Samples were cryosectioned at 10 µm thickness, and selected longitudinal spinal cord sections were positioned within the 10.5 × 22.5 mm capture areas of Xenium slides (10x Genomics). A custom 100-gene Xenium panel was designed to include markers of major spinal cord cell populations identified from the snRNA-seq dataset together with genes representing biological processes implicated in spinal cord regeneration (Table S2). In situ gene expression analysis was performed using the Xenium Analyzer and Xenium reagents according to the manufacturer’s instructions (CG000582, Rev A, 10x Genomics), and resulting spatial transcriptomic data were visualized and analyzed using Xenium Explorer v4 (10x Genomics).

### Spatial transcriptomic cell annotation and deconvolution

Cell type deconvolution of Xenium spatial transcriptomics data was performed using RCTD (Robust Cell Type Decomposition, spacexr v 2.2.1)(^75^). Xenium expression data were normalized and processed to obtain spatial coordinates and gene expression counts for individual cells by Seurat (v 5.5.0). The reference single-cell RNA-seq dataset was annotated using curated cell type labels, and raw gene counts together with cell type identities and UMI counts were used to construct the reference model. Spatial coordinates, expression counts, and total UMI counts from the Xenium dataset were used to generate the query object. RCTD was run in doublet mode to infer cell type compositions for each spatial cell. The resulting primary and secondary cell type assignments were integrated into the Xenium object.

### Data analysis and statistics

The number of animals analyzed for all quantifications (*n*) is indicated in the figure panels. Statistical analyses of experimental trials were performed using GraphPad Prism software (v10.4.1)

## Author Contributions

P.G. designed experimental strategy, generated mutant zebrafish lines, performed experiments, analyzed data and wrote the manuscript. S.B. designed experimental strategy, performed experiments using zebrafish and analyzed data. J.S. performed experiments using zebrafish. B.M and J.O. performed bioinformatics analysis. R.S. shared advice and technical expertise. K.D.P. designed experimental strategy, acquired funding, and reviewed and edited the manuscript. All authors commented on the manuscript.

## Acknowledgments

We thank Jim Burris, Mackenzie Klemek and Jacob Quijas at the Morgridge Institute for Research zebrafish facility and Lawrence Frauen, Colin Dolan, Kelly Scanlon, Makenna Powel and Maya Wiley at the Duke University zebrafish facility for animal care; the UW-Madison Biotechnology Center for their assistance with spatial transcriptomic experiments; and Ashley Rich, Rocky Diegmiller, and Valentina Cigliola for comments on the manuscripts.

## Data availability

The single-nuclei and spatial RNA sequencing datasets generated in this study have been deposited in the Gene Expression Omnibus (GEO) archive. The accession number for single-nucleus sequencing is GSE344595. The accession number for spatial RNA-seq data is GSE344338. Unique biological materials are available from the corresponding author upon request. All data files will be available at Dryad upon final acceptance for publication.

