## Supplementary Information for "Microenvironmental and transcriptional determinants of the robustness of spinal cord regeneration in zebrafish"

**Supplementary Information, includes 6 figures and 4 tables**

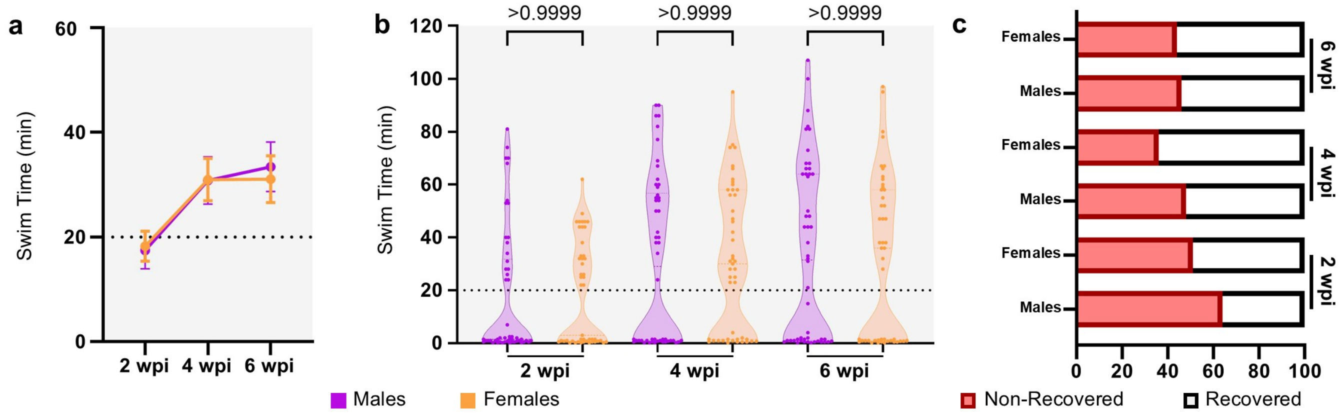


**Figure S1. Sex does not significantly influence functional recovery after spinal cord injury**

**A, B)** Swim capacity of males (magenta, n = 50) and females (orange, n =47) at increasing currents over a 6 week follow-up period after spinal cord transection. Data are shown as a line graph **(A)** at 2, 4 and 6 wpi and violin plot **(B)** at the similar timepoint. Each dot represents an individual fish. Dashed line indicates the 20 min acclimation period and serves as a threshold to identify non-recovered animals. Error bars in **(A)** indicate SEM. **(B)** Statistical comparison was performed using a Kruskal-Wallis multiple comparison test.

**C)** Recovered vs non-recovered proportion analysis between injured male and female adults at 2, 4 and 6 wpi. Recovery frequencies were comparable between male and female zebrafish across all cohorts (32/50 vs. 24/47, p = 0.22; 24/50 vs. 17/47, p = 0.29; 23/50 vs. 21/47, p = 1.00; Fisher's exact tests), indicating no significant effect of sex on recovery outcome following spinal cord injury.

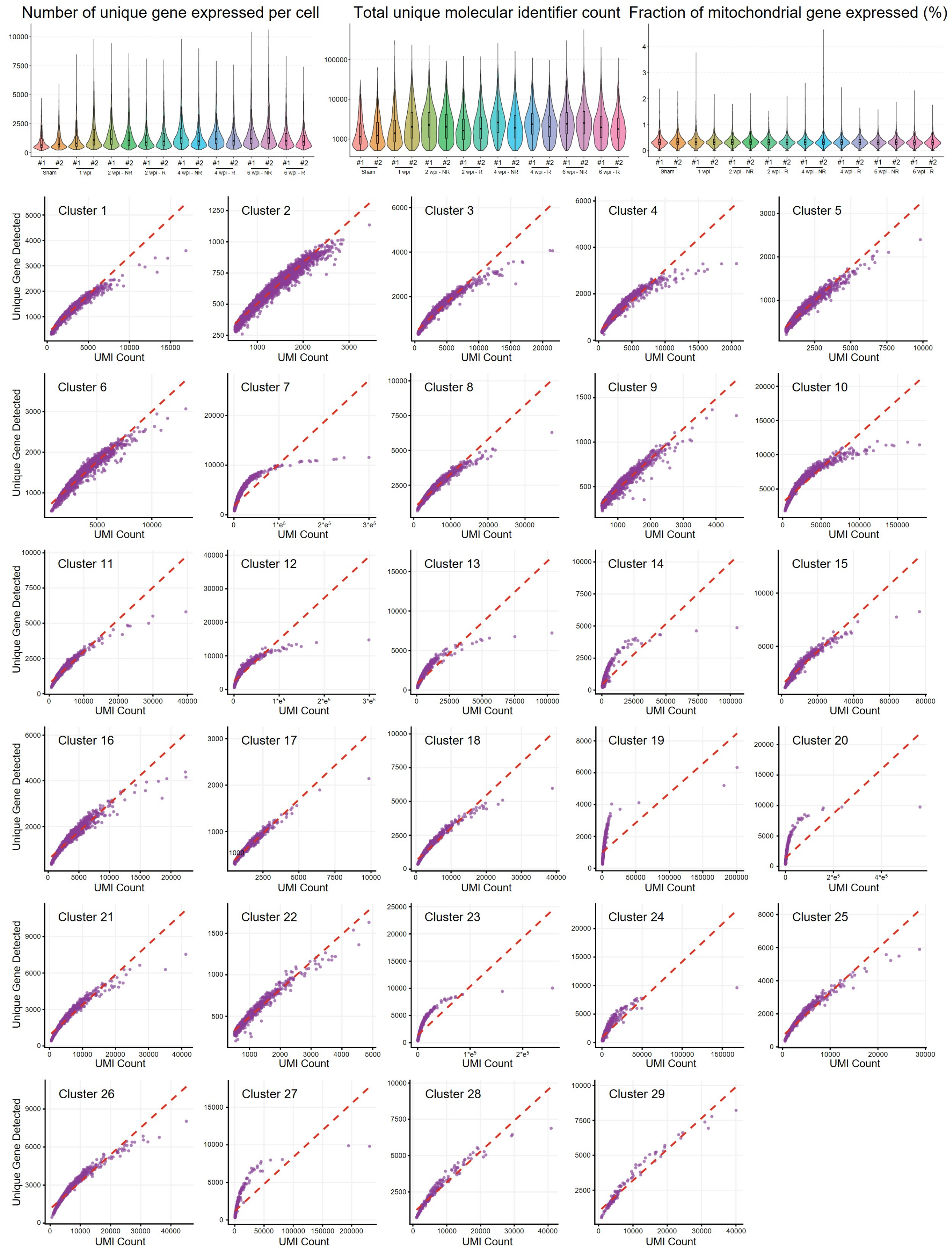
Number of unique gene expressed per cell
Total unique molecular identifier count
Fraction of mitochondrial gene expressed (%)
Sham
1 wpi
2 wpi - NR
2 wpi - R
4 wpi - NR
4 wpi - R
6 wpi - NR
6 wpi - R
Cluster 1
Cluster 2
Cluster 3
Cluster 4
Cluster 5
Cluster 6
Cluster 7
Cluster 8
Cluster 9
Cluster 10
Cluster 11
Cluster 12
Cluster 13
Cluster 14
Cluster 15
Cluster 16
Cluster 17
Cluster 18
Cluster 19
Cluster 20
Cluster 21
Cluster 22
Cluster 23
Cluster 24
Cluster 25
Cluster 26
Cluster 27
Cluster 28
Cluster 29
Unique Gene Detected
UMI Count

**Figure S2. Single-nuclei RNA-sequencing quality control across conditions and biological replicates**

**A-C)** Violin plots showing the distribution of cells based on their quality control metrics grouped by both biological replicates and conditions. Distributions of cells that were retained after low-quality removal are shown. The total number of unique genes found per cell **(A)**, the number of unique molecular identifiers (UMI) per cell **(B)**, and the proportion of UMIs that map to mitochondrial protein encoding genes per cell **(C)** are shown.

**D)** Scatter plots showing the distribution of nuclei within each cluster based on their total number of unique genes and their total number of UMIs.

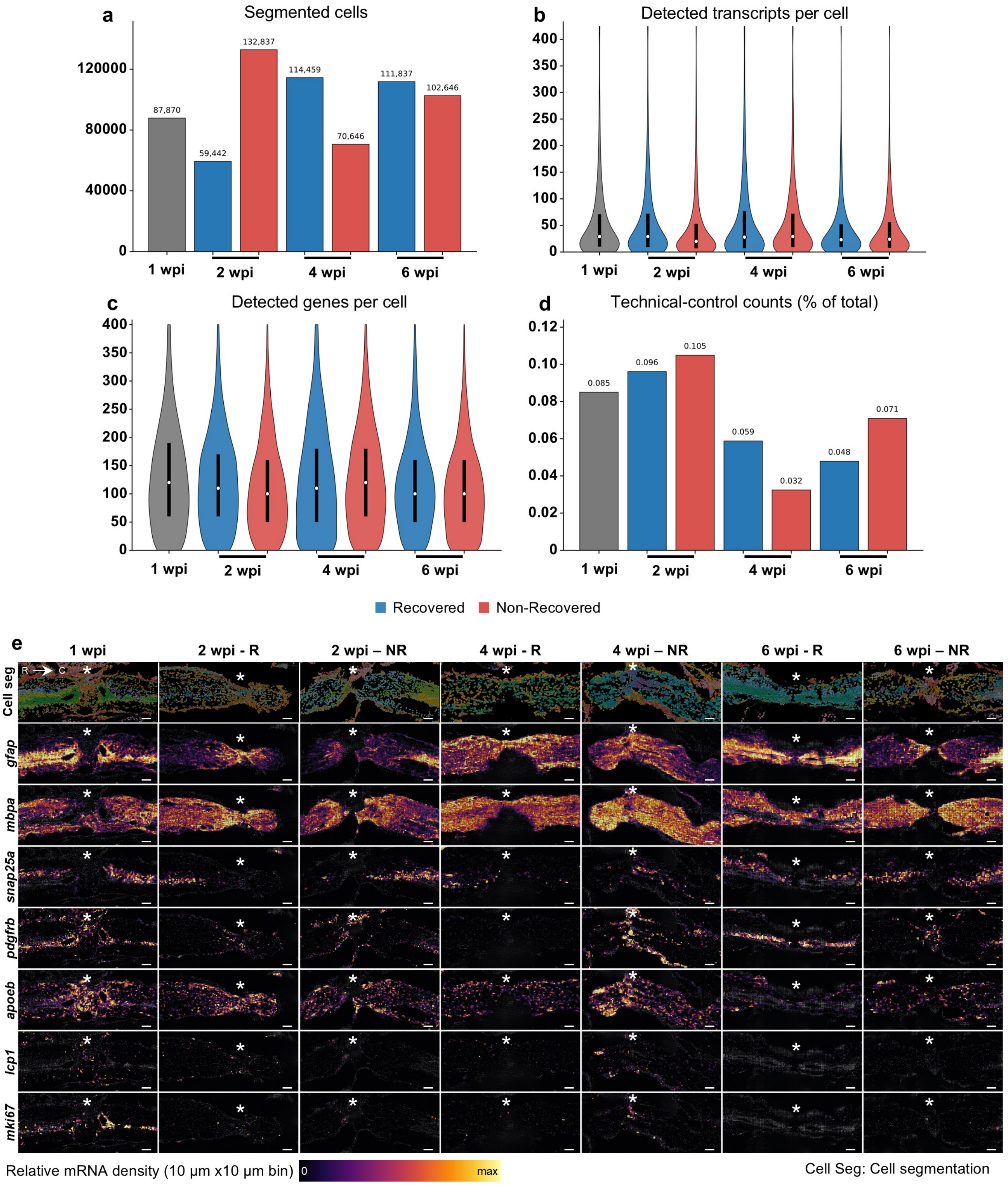

a
Segmented cells
120000
80000
40000
0
87,870
59,442
132,837
114,459
70,646
111,837
102,646
1 wpi
2 wpi
4 wpi
6 wpi
b
Detected transcripts per cell
400
350
300
250
200
150
100
50
0
c
Detected genes per cell
d
Technical-control counts (% of total)
0.12
0.10
0.08
0.06
0.04
0.02
0
0.085
0.096
0.105
0.059
0.032
0.048
0.071
Recovered
Non-Recovered
e
1 wpi
2 wpi - R
2 wpi – NR
4 wpi - R
4 wpi – NR
6 wpi - R
6 wpi – NR
Cell seg
R
C
gfap
mbpa
snap25a
pdgfrb
apoeb
lcp1
mki67
Relative mRNA density (10 µm x10 µm bin)
0
max
Cell Seg: Cell segmentation

**Figure S3. Quality control and spatial validation of the Xenium transcriptomic dataset**

**A)** Number of segmented cells identified in each Xenium dataset after pooling biological replicates for each experimental condition. Cell segmentation was performed using the Xenium analysis pipeline.

**B)** Distribution of detected transcript molecules per segmented cell across Xenium datasets. Violin plots represent the distribution of all segmented cells; white dots indicate the median and black bars the interquartile range.

**C)** Distribution of detected genes per segmented cell across Xenium datasets. Violin plots represent the distribution of all segmented cells; white dots indicate the median and black bars the interquartile range.

**D)** Fraction of transcripts assigned to Xenium technical control probes for each experimental condition, demonstrating low background signal across all datasets.

**E)** Density map expression of markers identifying most abundant cell types detected using 10x Xenium. Representative longitudinal sections of adult zebrafish spinal cord tissue during successful and failing regeneration (from 1 to 6 wpi) are shown. From top to bottom: glial cells (*gfap*), oligodendrocytes (*mbpa*), neurons (*snap25a*), fibroblasts (*pdgfrb*), microglia (*apoeb*), leukocytes (*lcp1*) and cycling cells (*mKI67*) are represented. Expression is shown as a relative RNA density heatmap (black to yellow) on 10 µm*10 µm bins. White asterisks indicate location of spinal cord transection site. Scale bars, 100 µm.

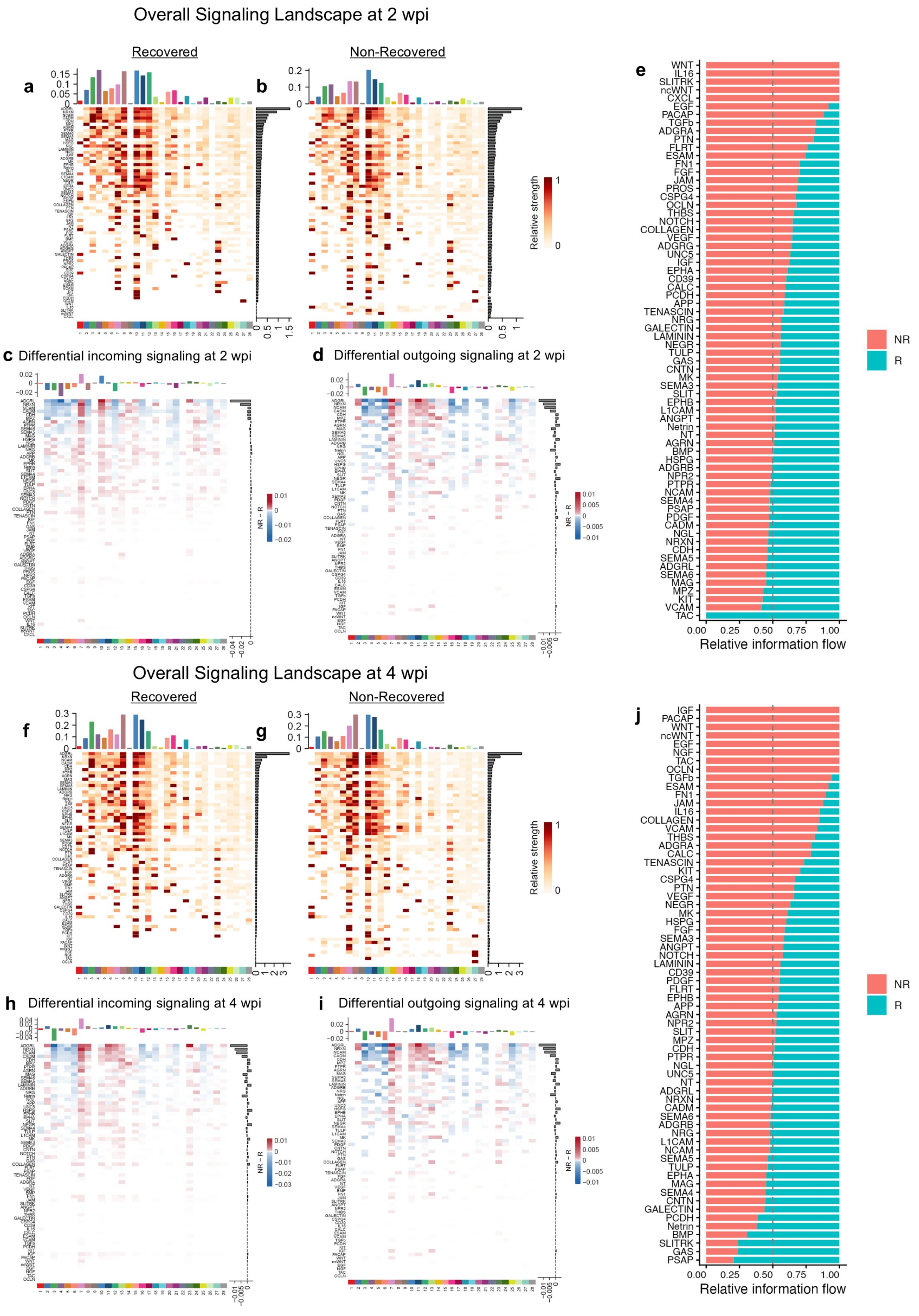

Overall Signaling Landscape at 2 wpi
Recovered
Non-Recovered
a
b
Relative strength
c Differential incoming signaling at 2 wpi
d Differential outgoing signaling at 2 wpi
NR - R
e
WNT
IL16
SLITRK
ncWNT
CXCL
EGF
PACAP
TGFb
ADGRA
PTN
FLRT
ESAM
FN1
FGF
JAM
PROS
CSPG4
OCLN
THBS
NOTCH
COLLAGEN
VEGF
ADGRG
UNC5
IGF
EPHA
CD39
CALC
PCDH
APP
TENASCIN
NRG
GALECTIN
LAMININ
NEGR
TULP
GAS
CNTN
MK
SEMA3
SLIT
EPHB
L1CAM
ANGPT
Netrin
NT
AGRN
BMP
HSPG
ADGRB
NPR2
PTPR
NCAM
SEMA4
PSAP
PDGF
CADM
NGL
NRXN
CDH
SEMA5
ADGRL
SEMA6
MAG
MPZ
KIT
VCAM
TAC
0.00 0.25 0.50 0.75 1.00
Relative information flow
NR
R
Overall Signaling Landscape at 4 wpi
Recovered
Non-Recovered
f
g
Relative strength
h Differential incoming signaling at 4 wpi
i Differential outgoing signaling at 4 wpi
NR - R
j
IGF
PACAP
WNT
ncWNT
EGF
NGF
TAC
OCLN
TGFb
ESAM
FN1
JAM
IL16
COLLAGEN
VCAM
THBS
ADGRA
CALC
TENASCIN
KIT
CSPG4
PTN
VEGF
NEGR
MK
HSPG
FGF
SEMA3
ANGPT
NOTCH
LAMININ
CD39
PDGF
FLRT
EPHB
APP
AGRN
NPR2
SLIT
MPZ
CDH
PTPR
NGL
UNC5
NT
ADGRL
NRXN
CADM
SEMA6
ADGRB
NRG
L1CAM
NCAM
SEMA5
TULP
EPHA
MAG
SEMA4
CNTN
GALECTIN
PCDH
Netrin
BMP
SLITRK
GAS
PSAP
0.00 0.25 0.50 0.75 1.00
Relative information flow
NR
R

**Figure S4. CellChat analysis reveals divergent intercellular signaling during successful and failed spinal cord regeneration**

**A, B)** Heatmaps summarizing the contribution of each cell population to significantly enriched signaling pathways inferred by CellChat in recovered (R) **(A)** and non-recovered (NR) **(B)** spinal cords at 2 wpi. Color intensity represents the relative contribution of each cell cluster to the inferred signaling pathway.

**C, D)** Differential signaling analysis comparing recovered and non-recovered animals. Heatmaps display changes in incoming **(C)** and outgoing **(D)** signaling strength for each cell population across all significant pathways at 2 wpi. Red indicates signaling enriched in non-recovered animals, whereas blue indicates signaling enriched in recovered animals.

**E)** Relative information flow of significant signaling pathways ranked according to their overall activity at 2 wpi. Bar coloration represents the contribution of recovered-associated cells (blue) and non-recovered-associated cells (red) to the associated signaling pathway, highlighting signaling programs preferentially associated with regenerative success or spontaneous regenerative failure.

**F, G)** Heatmaps summarizing the contribution of each cell population to significantly enriched signaling pathways inferred by CellChat in recovered (R) **(F)** and non-recovered (NR) **(G)** spinal cords at 4 wpi. Color intensity represents the relative contribution of each cell cluster to the inferred signaling pathway.

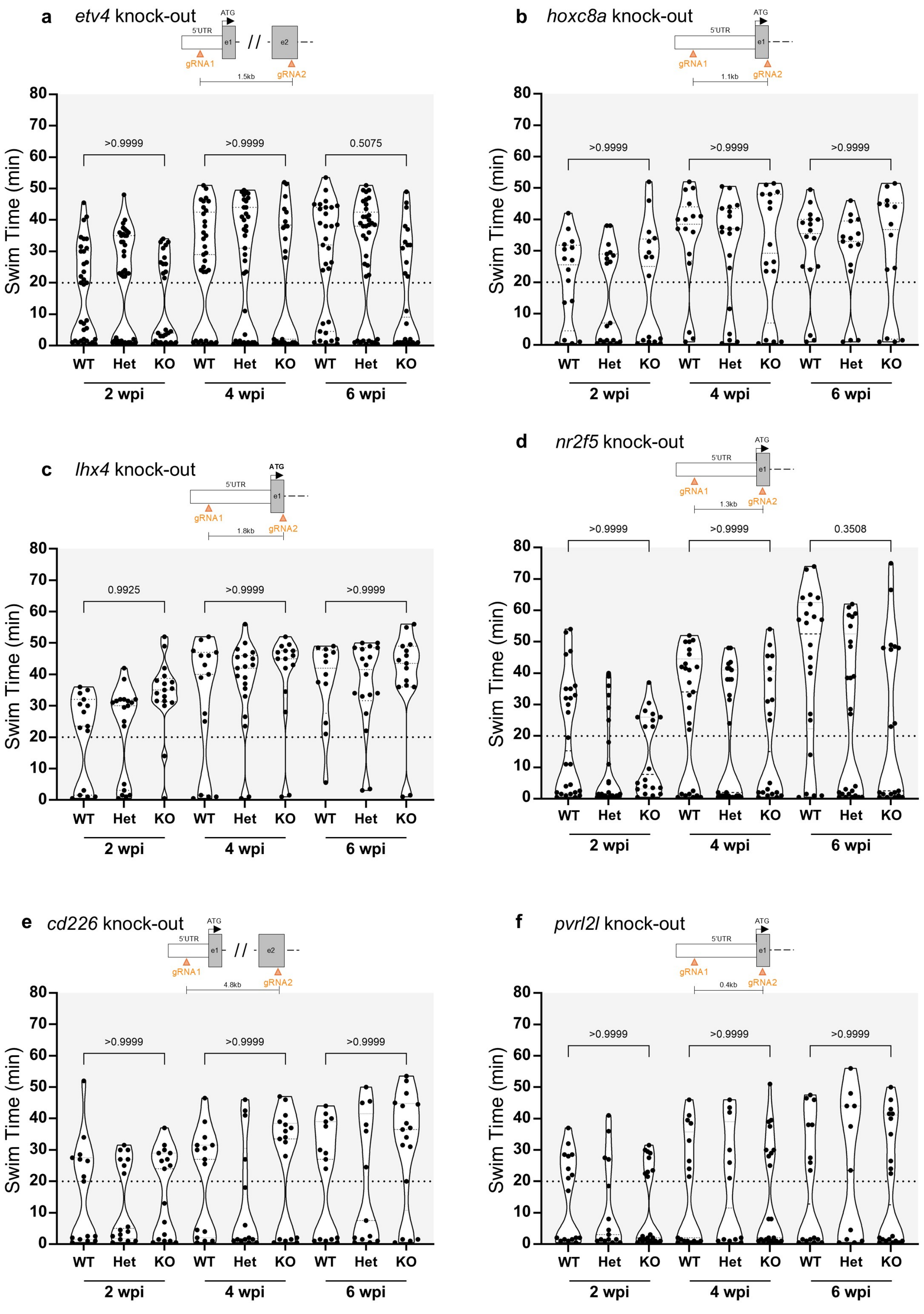
a etv4 knock-out
ATG
5'UTR
e1
e2
gRNA1
gRNA2
1.5kb
b hoxc8a knock-out
1.1kb
c lhx4 knock-out
1.8kb
d nr2f5 knock-out
1.3kb
e cd226 knock-out
4.8kb
f pvrl2l knock-out
0.4kb
Swim Time (min)
WT
Het
KO
2 wpi
4 wpi
6 wpi
>0.9999
0.5075
0.9925
0.3508

**Figure S5. Stable mutant lines for additional candidate regulators show no impairment in functional recovery**

**A-F)** Mutation schematic depicting the deleted region and swim capacity of stable mutant lines generated in this study over a 6 week periods following spinal cord surgery. Results are shown for *etv4* **(A)**, *hoxc8a* **(B)**, *lhx4* **(C)**, *nr2f5* **(D)**, *cd226* **(E)** and *pvrl2l* **(F)**. Data are shown as violin plots with each dot representing an individual animal. Dashed line indicates the 20 min acclimation period and serves as a threshold to identify non-recovered animals. Two-way repeated-measures ANOVA tests with Holm-Šidák correction were used to calculate exact p-values; only p-values corresponding to WT vs KO at 2, 4 and 6 wpi are shown. No significant differences were found.

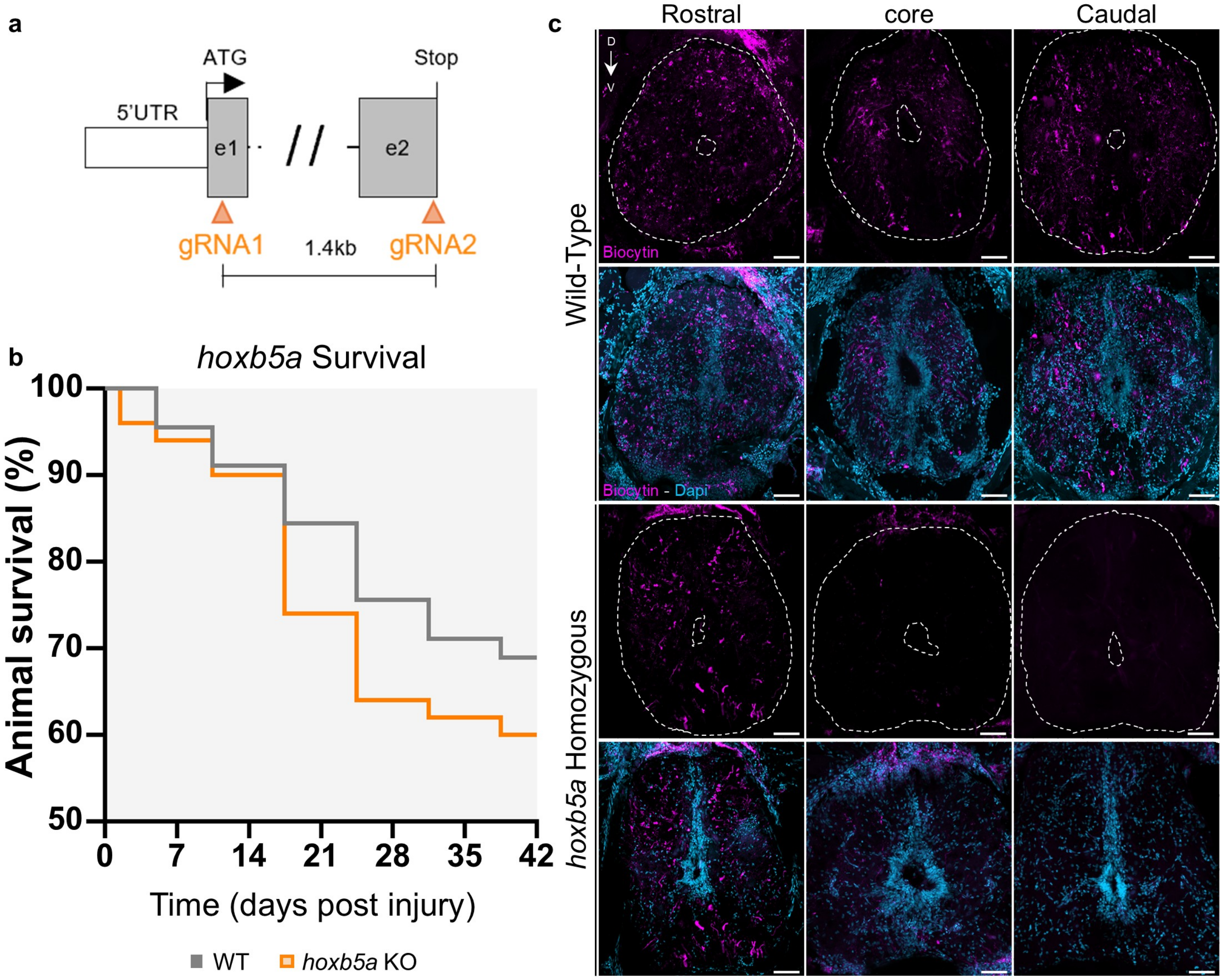
a
ATG
Stop
5'UTR
e1
e2
gRNA1
1.4kb
gRNA2
b
hoxb5a Survival
Animal survival (%)
100
90
80
70
60
50
0
7
14
21
28
35
42
Time (days post injury)
WT
hoxb5a KO
c
Rostral
core
Caudal
D
V
Wild-Type
Biocytin
Biocytin - Dapi
hoxb5a Homozygous

**Figure S6. Generation and characterization of the stable *hoxb5a* mutant line**

**A)** Schematic representation depicting the exon regions deleted to generate the *hoxb5a* mutant line.

**B)** Survival curve of *hoxb5a* mutants after spinal cord surgery. Survival was monitored from the day of injury (0 dpi) to 6 wpi, and the percentage of animals remaining alive at each observation time is shown. Statistical comparisons between genotypes at each time point were performed using two-sided Fisher's exact tests with Holm correction for multiple comparisons. Initial cohort sizes were n = 45 (WT) and n = 50 (KO).

**C)** Transverse sections of WT and hoxb5a homozygous mutant spinal cords shown in Figure 6f, biocytin (magenta) and DAPI (light blue) staining. Scale bars: 50 µm.

### Supplementary Tables

**Table S1. Single-nucleus RNA-seq cluster markers and gene expression.** Marker genes and gene-expression values used to characterize and annotate the transcriptionally distinct cell populations identified in the adult zebrafish spinal cord snRNA-seq dataset.

**Table S2. Xenium probe panel.** Genes included in the custom 100-gene Xenium panel for targeted spatial transcriptomics, selected to represent major spinal cord cell identities and biological processes associated with spinal cord regeneration.

**Table S3. Differential expression and Gene Ontology analysis from T-cell and fibroblast data.** Differentially expressed genes and Gene Ontology Biological Process (GO-BP) enrichment results for T cells and fibroblasts comparing recovered, non-recovered and sham-injured spinal cords across regeneration time points.

**Table S4. Gene regulatory network analysis.** Summary of CellOracle-inferred gene regulatory networks, including condition-specific GRN rewiring, transcription factor centrality and differential centrality, transcription factor family-level analyses, and predicted *hoxb5a* and *foxp4* regulons in recovered and non-recovered spinal cords.
